# New flat embedding method for transmission electron microscopy reveals an unknown mechanism of tetracycline

**DOI:** 10.1101/820191

**Authors:** Michaela Wenzel, Marien P. Dekker, Biwen Wang, Maroeska J. Burggraaf, Wilbert Bitter, Jan R. T. van Weering, Leendert W. Hamoen

## Abstract

Transmission electron microscopy (TEM) is an important imaging technique in bacterial research and requires ultrathin sectioning of resin embedding of cell pellets. This method consumes milli- to deciliters of culture and results in sections of randomly orientated cells. For rod-shaped bacteria, this makes it exceedingly difficult to find longitudinally cut cells, which precludes large-scale quantification of morphological phenotypes. Here, we describe a new fixation method using either thin agarose layers or carbon-coated glass surfaces that enables flat embedding of bacteria. This technique allows for the observation of thousands of longitudinally cut rod-shaped cells per single section and requires only microliter culture volumes. We successfully applied this technique to Gram-positive *Bacillus subtilis*, Gram-negative *Escherichia coli*, the tuberculosis vaccine strain *Mycobacterium bovis* BCG, and the cell wall-lacking mycoplasma *Acholeplasma laidlawii*. To assess the potential of the technique to quantify morphological phenotypes, we examined cellular changes induced by a panel of different antibiotics. Surprisingly, we found that the ribosome inhibitor tetracycline causes significant deformations of the cell membrane. Further investigations showed that the presence of tetracycline in the cell membrane changes membrane organization and affects the peripheral membrane proteins MinD, MinC, and MreB, which are important for regulation of cell division and elongation. Importantly, we could show that this effect is not the result of ribosome inhibition but is a secondary antibacterial activity of tetracycline that has defied discovery for more than 50 years.

**Significance:** Bacterial antibiotic resistance is a serious public health problem and novel antibiotics are urgently needed. Before a new antibiotic can be brought to the clinic, its antibacterial mechanism needs to be elucidated. Transmission electron microscopy is an important tool to investigate these mechanisms. We developed a flat embedding method that enables examination of many more bacterial cells than classical protocols, enabling large-scale quantification of phenotypic changes. Flat embedding can be adapted to most growth conditions and microbial species and can be employed in a wide variety of microbiological research fields. Using this technique, we show that even well-established antibiotics like tetracycline can have unknown additional antibacterial activities, demonstrating how flat embedding can contribute to finding new antibiotic mechanisms.

## Introduction

Transmission electron microscopy (TEM) is a powerful tool to examine the morphology and ultrastructure of bacterial cells. There are many bacterial embedding protocols for TEM (1–5), but the basic procedure, i.e. embedding of cell pellets as small nuggets into resin blocks, has not changed since the beginning of electron microscopy research on bacteria 60 years ago (4, 6, 7). This technique has two major shortcomings. Most importantly, it results in random orientations of cells in the ultrathin sections. This is a critical limitation when examining rod-shaped and other non-coccoid bacterial species, since the vast majority of cells are randomly cross-sectioned, and the number of complete longitudinally cut cells is generally so low that robust quantification and population-wide studies are not feasible. Another limitation is that acquiring a concentrated cell pellet often requires relatively large culture volumes typically in the range of 10 to 50 ml (6, 8, 9). This can be problematic when studying the mode of action of experimental antimicrobial compounds, whose synthesis or purification is laborious and expensive.

We have addressed these problems by developing a novel embedding technique that enables observation of a large number of cells oriented in one plane by immobilizing bacterial samples on a flat surface of either agarose or glass. This relatively simple method does not require any expensive equipment and can be adapted for any microorganism. We have successfully used this method with the Gram-positive bacterium *Bacillus subtilis*, the Gram-negative bacterium *Escherichia coli*, the tuberculosis vaccine strain *Mycobacterium bovis* BCG, and the cell wall-less mycoplasma species *Acholeplasma laidlawii*. This flat embedding technique allowed the quantification of morphological changes in bacteria treated with different antibiotics. This led to the surprising discovery that the well-known ribosome inhibitor tetracycline does not only block translation but also directly disturbs the bacterial cell membrane. This additional mechanism of action has remained hidden for over 50 years despite the fact that tetracyclines are one of the most commonly used antibiotic groups in both human and veterinary medicine (10).

## Results

### Alignment of cells on agarose

Light microscopy studies of bacteria commonly use thin agarose layers to immobilize cells (11, 12). If done correctly, these cells are well-aligned in a single plane, allowing large-scale quantification of phenotypic changes. We wondered whether this immobilization technique could be adapted for TEM embedding, which would solve the issue of randomly sectioned bacteria and at the same time drastically reduce the required sample volume. Using rod-shaped *B. subtilis* cells as model sample, we tested different conditions, eventually resulting in the following flat embedding procedure. As little as 50-150 µl of logarithmically growing (OD_600_ = 0.4) *B. subtilis* culture was pelleted, resuspended in 5-15 µl medium and spotted on a thin, flat layer of 1.5% agarose (Figure 1A, Supplementary Movies 1 and 2). After evaporation of excess liquid, the immobilized cells were subjected to a standard sequence of fixation, staining, dehydration, and finally resin embedding, resulting in an EPON disc carrying the flat embedded bacteria (Supplementary Figure 1). Some cells were washed off during the procedure, but the majority remained attached to the agarose and was successfully embedded. As shown in Figure 1B, cells were generally well-aligned in the resulting ultrathin sections. Only 5 images of a single ultrathin section were sufficient to examine more than 900 individual fully longitudinally sectioned cells (5000x magnification). When we examined TEM pictures of bacteria prepared with the classical pellet embedding method, we found on average only 6 fully longitudinally sectioned bacteria per image (Figure 1B). Even filamentous cell division mutants, which normally pose a particular challenge for TEM, could be efficiently sectioned longitudinally using this new flat embedding protocol (Supplementary Figure 2).

**Figure 1:**
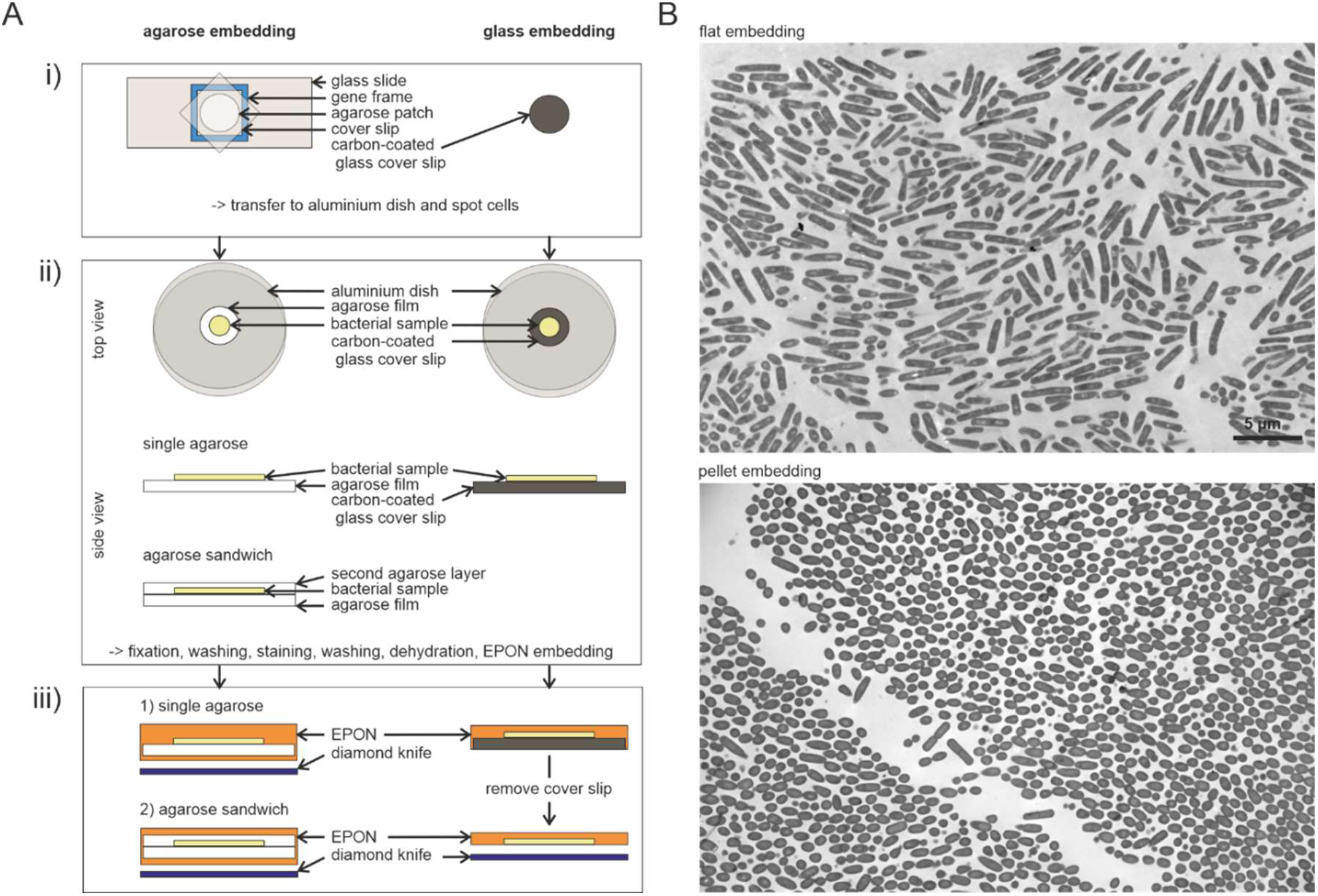
The flat embedding technique. (A) Schematic representation of the flat embedding work flow including embedding on a single layer of agarose, embedding in an agarose sandwich (for mycobacteria), and embedding on carbon-coated glass coverslips. (i) Preparation of the surface. Uniform thickness of the agarose film is ensured by using a gene frame as spacer. (ii) The agarose or glass surface is transferred to an aluminum dish and a small drop of cell sample is spotted on top of the surface and allowed to dry under a slight air flow. For the agarose sandwich approach, a second flat layer of agarose is added on top of the cells without using a gene frame. (iii) Samples after EPON embedding. For glass embedding, the glass cover slip is broken off the EPON disc prior to sectioning. (B) Overview pictures of *B. subtilis* 168 cells embedded on a single agarose layer (top) and as pellet (bottom) at 5000x magnification.

### Flat embedding applied to different bacteria

To examine whether flat embedding is applicable to a wider range of microorganisms, we tested bacterial species with different cell surface properties. *E. coli* was chosen as representative of Gram-negative bacteria, *M. bovis* BCG as representative of bacteria with a mycolic acid-containing outer membrane, and *A. laidlawii* as a cell wall-less mycoplasma species. Both *E. coli* and *A. laidlawii* were easy to embed on agarose (Figure 2A). However, *M. bovis* BCG was easily washed off the agarose surface during subsequent washing and fixation steps, resulting in only very few cells being left on the final sections. Typically, *M. bovis* BCG is grown in the presence of detergent (Tween 80) to reduce clumping and to facilitate microscopic observation of single cells (13, 14). However, the presence of detergent might reduce the mycobacterial capsule and affect cell morphology (13, 15–20), and we hypothesized that it might also affect the attachment of the cells to the agarose patch. However, growing *M. bovis* BCG without detergent did not improve attachment to the agarose surface. On the contrary, clumping cells detached even more readily and could not be embedded with this method. To overcome this problem, we developed an agarose sandwich approach. To this end, cells were covered with a second thin layer of agarose after spotting on the first flat agarose layer (Figure 1A, Supplementary Movie 2). Using this approach, we were able to easily embed both detergent-treated and detergent-free cultures of *M. bovis* (Figure 2B). Thus, flat agarose patches can be used to immobilize a wide variety of bacterial cells in a single plane for longitudinal TEM sectioning.

**Figure 2:**
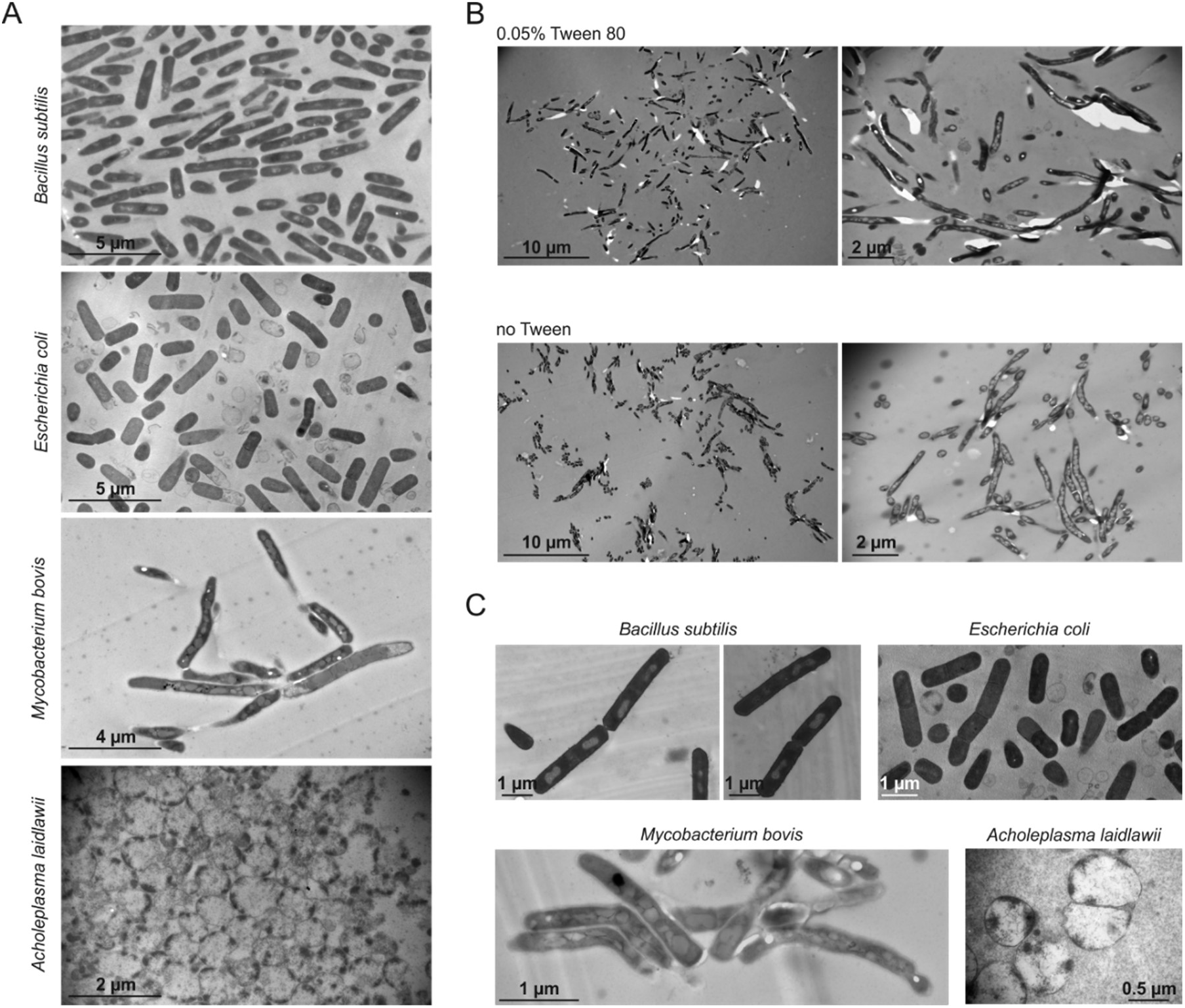
Transmission electron micrographs of flat-embedded *Bacillus subtilis, Escherichia coli, Mycobacterium bovis* BCG, and *Acholeplasma laidlawii.* (A) Bacteria embedded on a single layer of agarose. (B) Agarose sandwich approach to flat embedding of mycobacteria. *M. bovis* BCG was grown in the presence or absence of 0.05% Tween 80. (C) Bacteria embedded on carbon-coated coverslips.

### Flat embedding on carbon-coated glass surfaces

While flat embedding on agarose was easy and straight-forward, it can be time-consuming to find the perfect plane during ultrathin-sectioning. To facilitate this step, we developed an alternative embedding method by spotting cells onto a carbon-coated glass cover slip (Figure 1A). The carbon film was applied to better visualize the bacteria during sectioning. After embedding, the glass was removed from the polymerized resin, leaving the cells very close to the surface of the EPON disc. This, and the easy localization of the cells due to the contrast of the dark carbon film greatly facilitated finding the right section plane. Since only cells and no agarose patches have to be dehydrated in this protocol, it is significantly faster at the embedding stage as well. It also eliminates the risk of artefacts caused by insufficient dehydration of the agarose film, which can complicate sectioning and produces ‘waves’ in the sections. As shown in Figure 2C, embedding on glass worked for all tested species and resulted in flat, clean, and nicely sectioned samples. However, cells detached easier from glass than from the agarose surface, resulting in less cells in the final sections.

### Antibiotic mode of action studies

Our new flat embedding method enables a quantitative approach to monitor antibiotic-induced cell damage using TEM. To demonstrate this, we counted antibiotic-induced phenotypic changes in at least 100 *B. subtilis* cells caused by a panel of well-characterized antimicrobial compounds, including vancomycin, ampicillin, daptomycin, MP196, nitrofurantoin and tetracycline. Concentrations were used that clearly reduced the growth rate without causing extensive cell lysis (Supplementary Table 1, Supplementary Figure 3). After incubation of cultures with the selected antibiotic concentrations for 30 min, samples were embedded using the single layer agarose approach. Typical examples of cells exhibiting cellular aberrations that are characteristic for the individual antibiotics are shown in Supplementary Figure 4. While ampicillin, daptomycin and MP196 caused the expected phenotype (see legend of Supplementary Figure 4 for details), vancomycin, nitrofurantoin and tetracycline displayed unexpected phenotypes and were chosen for further analysis. Vancomycin, a last line of defense antibiotic for systemic Gram-positive infections, binds to the peptidoglycan precursor molecule lipid II (21). Cells treated with this antibiotic showed characteristic cell envelope lesions that are indicative of aberrant cell wall synthesis. However, only in 32% of cells vancomycin-induced lesions occurred at the cell periphery, while in 44% of the cells, they were located at cell poles and 22% at cell division septa (Figure 3C). The locally increased concentration of lipid II at developing septa (22) might explain the higher proportion of lesions occurring at new and old cell division sites, i.e. cell poles.

**Figure 3:**
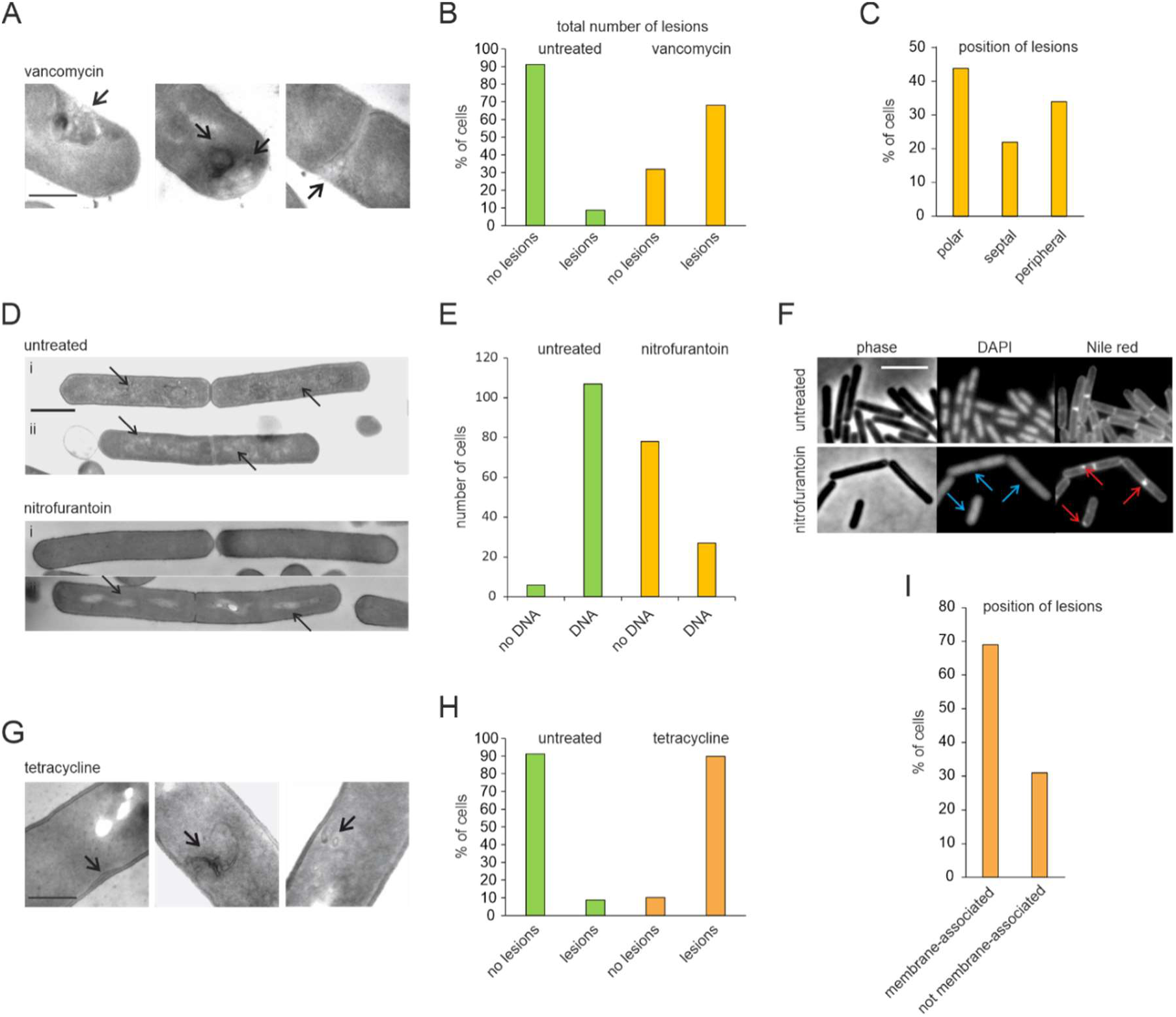
Quantification of antibiotic-induced lesions based on electron micrographs. Cells were quantified from electron micrographs at 8000 to 15000x magnification. A minimum of 100 cells was examined per condition. (A) TEM images showing cell wall damage caused by vancomycin, which occurred either at the periphery (left panel), the poles (middle panel), or the septum (right panel). (B) Quantification of the total number of lesions caused by vancomycin. (C) Position of cell wall lesions in vancomycin-treated cells. (D) TEM images showing deterioration of nucleoids caused by nitrofurantoin. Electron micrographs show two different examples of untreated control cells (upper panels; i and ii: healthy control cells with visible nucleoids) and nitrofurantoin-treated cells (lower panels; i: cell without DNA, ii: cell with deteriorated nucleoid). Arrows indicate nucleoid structures. Scale bar 1 µm. (E) Quantification of cells devoid of visible nucleoids in electron micrographs. Note that all nitrofurantoin-treated cells with visible DNA structures displayed a deteriorated nucleoid as shown in A (lower panel ii). (F) Fluorescence light microscopy images of *B. subtilis* stained with the DNA dye DAPI and the membrane dye Nile red. Blue arrows indicate diffuse DNA stain. Red arrows indicate membrane patches. Cells were treated with 4x MIC of nitrofurantoin for 30 min. Scale bar 3 µm. (G) TEM images showing lesions caused by tetracycline that are either clearly membrane-associated (left and middle panels) or not visibly membrane-associated (right panel). Scale bar 500 nm. (H) Quantification of total lesions in cells treated with tetracycline. (I) Quantification of different types of membrane lesions caused by tetracycline.

Nitrofurantoin is an antibiotic that is widely used against urinary tract infections since 1953, but its mechanism is still not fully understood. It is thought to damage DNA, RNA, proteins, and other macromolecules by a mechanism involving oxidative damage (23). Interestingly, in our TEM images 74% of cells treated with nitrofurantoin seemed to entirely lack a nucleoid, whereas the other 26% showed condensed remnants of chromosomes (Figure 3D,E). To confirm this finding, cells were stained with the fluorescent DNA dye DAPI and examined by fluorescence light microscopy. Already after 5 min of treatment cells stared to show condensed nucleoids (Supplementary Figure 5) and after 30 min the DAPI signal became completely diffuse (Figure 3F). The TEM images also showed accumulation of small membrane vesicles (Supplementary Figure 4) and 50% of cells also showed fluorescent membrane patches when stained with the membrane dye Nile red (Figure 3F). Both DNA and lipids are sensitive to oxidative damage (24–26) and our results corroborate the current model of nitrofurantoin action.

The commonly used antibiotic tetracycline is known to inhibit protein biosynthesis by blocking binding of the aminoacyl-tRNA to the ribosome (10, 27). Interestingly, 90% of tetracycline-treated cells exhibited cellular lesions in the TEM images reminiscent of membrane invaginations (Figure 3G,H). The majority of these (69%) were visibly membrane-associated (Figure 3I). These results may suggest that tetracycline does not only target the ribosome but also affects the bacterial cell membrane.

### Tetracycline is a membrane-active compound

To investigate the effect of tetracycline on bacterial cell membranes in more detail, we tested whether we could observe membrane deformations with fluorescence microscopy using the membrane dye Nile red. As shown in Figure 4A, tetracycline indeed caused aberrant, highly fluorescent membrane patches in 93% of cells (Supplementary Figure 6). We were able to localize the antibiotic directly due to its green autofluorescence, which appeared to overlap with Nile-red stained membrane foci (Figure 4A). This irregular green fluorescence membrane staining was also observed in cells that were not stained with Nile red (Supplementary Figure 7), indicating that it is not a fluorescence bleed-through artifact from the membrane dye.

**Figure 4:**
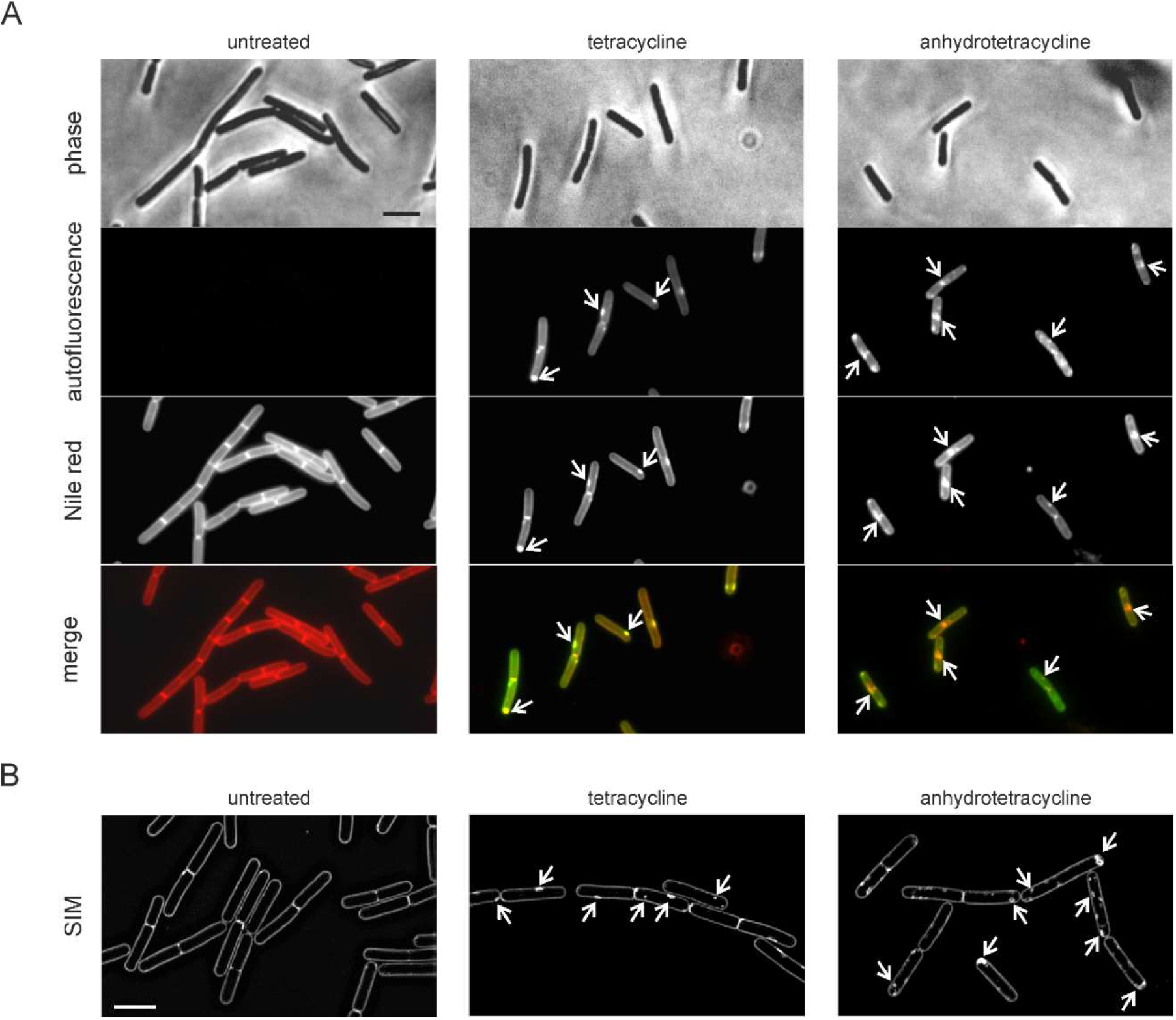
Tetracycline targets the cytoplasmic membrane. (A) Fluorescence microscopy images of cells treated with either tetracycline or anhydrotetracycline for 30 min. Both compounds display green autofluorescence allowing direct localization of the compound. Cell membranes were stained with Nile red. Arrows indicate fluorescent membrane patches coinciding with accumulation of the respective antibiotic. See Supplementary Figure 6 for quantification. (B) SIM microscopy images of cells treated with either tetracycline or anhydrotetracycline for 30 min. Membranes were stained with Nile red. Arrows indicate membrane staples or invaginations. Scale bars 2 µm.

The TEM images suggested that the highly fluorescent Nile red patches are likely caused by the accumulation of extra membrane material due to membrane invaginations (28–30) (Figure 3A, Supplementary Figures 4). To confirm this, we increased the fluorescence microscopy resolution by employing Structured Illumination Microscopy (SIM). This revealed clear membrane invaginations after treatment with tetracycline (Figure 4B).

The tetracycline analogue anhydrotetracycline is broadly applied in molecular genetics as inducer of Tet repressor-based gene expression systems (31, 32), since it is widely believed not to inhibit translation or bacterial growth (33). Interestingly, incubation of *B. subtilis* cells with anhydrotetracycline also caused fluorescent Nile red foci that appear to be caused by membrane invaginations (Figure 4A-B, Supplementary Figure 6,7). These results confirmed our observations by TEM (Supplementary Figure 4,8), suggesting that tetracycline affects the bacterial cell membrane independently from the inhibition of protein translation.

### Tetracycline affects membrane protein localization

To test whether tetracycline functionally disturbs the cell membrane, we examined the localization of three peripheral membrane proteins that are known to be affected by membrane depolarization, MinC, MinD and MreB (34, 35). MinD interacts with MinC to form a complex that inhibits initiation of cell division at the cell poles (36). Using a strain that expresses a GFP fusion to MinD and an mcherry fusion to MinC (37), we observed that the localization of both proteins was severely disturbed by both tetracycline and anhydrotetracycline after only 5 min of treatment (Figure 5A). MreB is an actin homologue that forms dynamic polymers along the lateral membrane and coordinates lateral cell wall synthesis (38). Tetracycline slightly affected localization of MreB and caused gaps in the normally regular localization pattern of this protein (Figure 5B). Anhydrotetracycline caused a much more dramatic effect and completely delocalized MreB resulting in diffuse fluorescence signal and large local clusters (Figure 5B).

**Figure 5:**
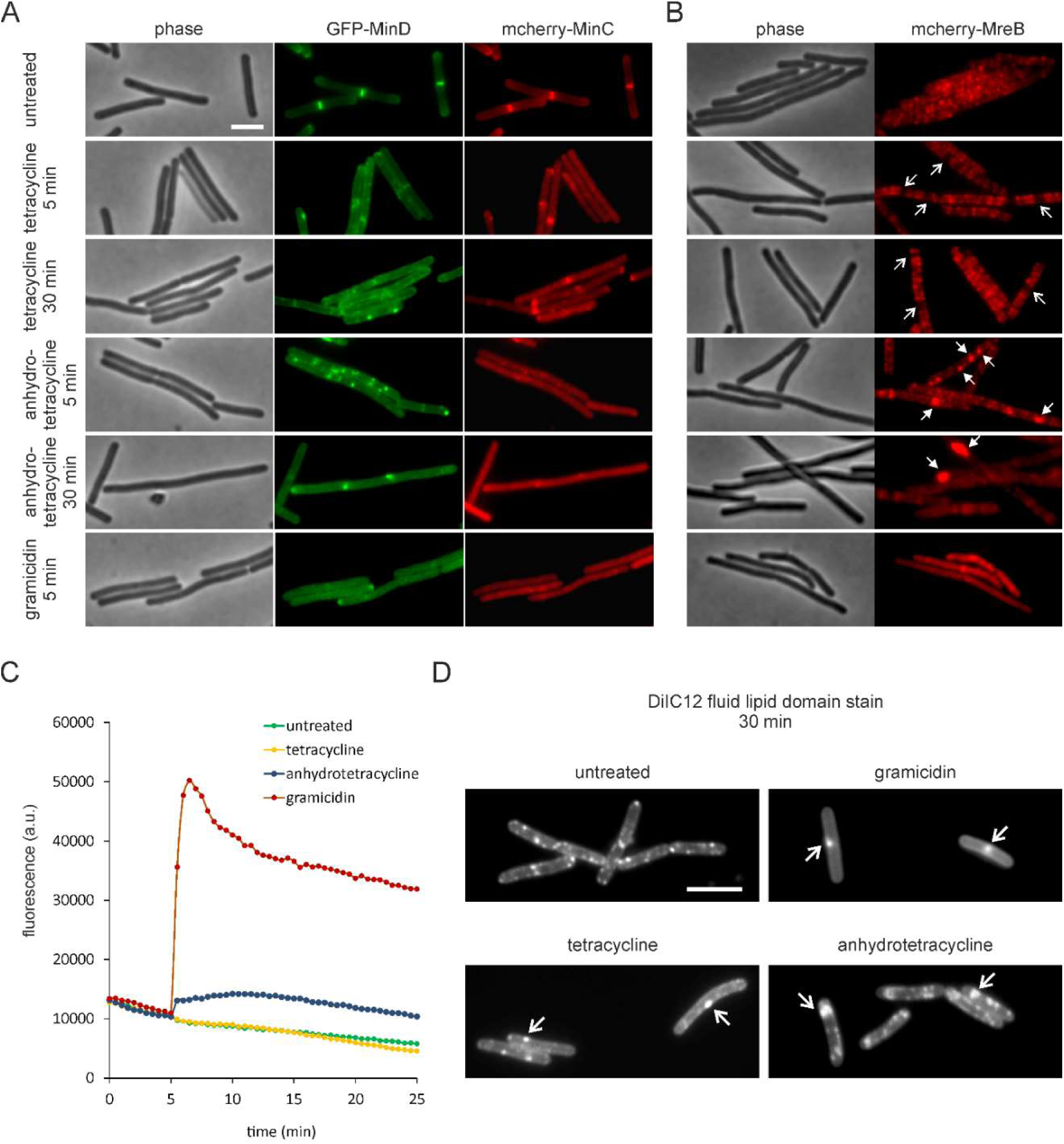
Tetracyclines delocalize membrane proteins. Effects of tetracycline (2 µg/ml), anhydrotetracycline (2 µg/ml), and gramicidin (1 µg/ml, positive control) on *B. subtilis* LB318 (168 *amyE::spc mgfp-minD aprE::cat mcherry-minC*) (A) and TNVS205 (168 *aprE::cat mcherry-mreB*) (B). (C) Effects of tetracycline (2 µg/ml), anhydrotetracycline (2 µg/ml), and gramicidin (1 µg/ml, positive control) on the membrane potential of *B. subtilis* 168 measured with DiSC(3)5. An exemplary graph out of three biological replicates is shown. (D) Effects of tetracycline (2 µg/ml), anhydrotetracycline (2 µg/ml), and gramicidin (1 µg/ml, positive control) on fluid membrane microdomains of *B. subtilis 168* cells stained with the fluid lipid domain dye DiIC12. Scale bars 2 µm.

### Tetracycline does not dissipate the membrane potential

Since the localization of MinC, MinD, and MreB depends on the membrane potential, we wondered whether tetracycline depolarizes the cell membrane. This was tested using the voltage-sensitive probe DiSC(3)5, which accumulates in bacterial cells in a membrane potential-dependent manner (11). As shown in Figure 5C, no depolarization of the cell membrane was observed, even after 30 min of incubation. Anhydrotetracycline caused only a slight membrane depolarization. The latter effect was not due to the presence of a subset of cells that had lost their membrane potential (Supplementary Figures 9–11). These results show tetracyclines disturb the bacterial cell membrane by a mechanism that is unrelated to membrane permeabilization.

### Tetracycline disturbs membrane organization

Bacteria can contain specific membrane regions of increased fluidity called RIFs (38, 39). RIFs contain fluidizing lipid species, e.g. with short, branched or unsaturated fatty acid chains. Since insertion of a membrane anchor into a lipid bilayer is facilitated in a more fluid environment, RIFs are enriched in certain peripheral membrane proteins (38, 40, 41). MreB is associated with RIFs (38) and its observed delocalization could be an indication that tetracycline affects these lipid domains. RIFs can be visualized with the fluidity-sensitive dye DiIC12 (12, 38). As shown in Figure 5D, tetracycline, and especially anhydrotetracycline, affected the formation of RIFS in a different manner than the membrane-depolarizing peptide gramicidin. Thus, the delocalization of MinCD and MreB is a consequence of the distortion of lipid organization by the tetracyclines, and not because of membrane depolarization.

### Membrane activity is independent of ribosome inhibition

Several observations suggested that tetracycline directly targets the bacterial cell membrane independently from inhibition of ribosomes. Firstly, the antibiotic visibly accumulated in the membrane lesions observed by fluorescent microscopy (Figure 4A). Secondly, the localization of membrane proteins was affected after a short treatment time (Figure 5). Thirdly, membrane deformations caused by tetracycline are largely similar of those of anhydrotetracycline (Figures 4 and 5, Supplementary Figures 4,6–8). As an additional control, we tested the effects of the translation inhibitors chloramphenicol and kanamycin on membrane organization. Neither chloramphenicol nor kanamycin caused membrane invaginations, affected the localization of MinCD and MreB, or affected RIFS (Supplementary Figure 12–15).

Finally, we analyzed two different tetracycline-resistant strains, the *tet-4* point mutation in the ribosomal protein S10, which reduces the tetracycline sensitivity of the ribosome (42, 43), and a strain containing the *tetL* resistance cassette, which encodes the TetA tetracycline transporter and confers high-level tetracycline resistance (44). If the effects of tetracycline on the membrane are a consequence of ribosome inhibition, they should be absent in both the *tet-4* and *tetL* mutant. As shown in Figure 6, membrane distortions were still clearly visible in the *tet-4* mutant, indicating that the interaction of tetracycline with ribosomes is not required for its membrane activity. In contrast, the *tetL* mutant showed no membrane lesions, which makes sense since TetA is an efflux pump that removes tetracycline from the membrane (44).

**Figure 6:**
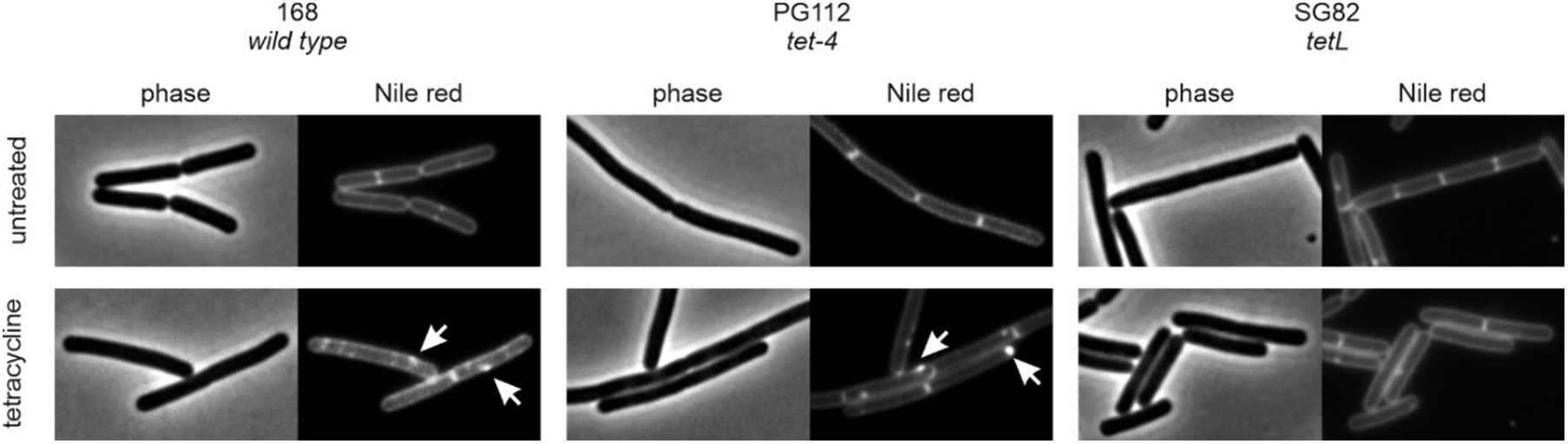
Tetracycline affects the membrane independently of ribosome inhibition. Strain PG112 carries the *tet-4* mutation, a point mutation in the ribosomal protein S10 that renders ribosomes insensitive to tetracycline. Strain SG82 carries the *tetL* tetracycline efflux pump. Cells were treated with 1x MIC of tetracycline (16 µg/ml for PG112, 100 µg/ml for SG82) for 30 min prior to membrane staining with Nile red and microscopy. Scale bar 2 µm.

## Discussion

Here we described a new method for embedding bacterial cells in a single layer to facilitate observation by TEM. This technique enabled us to observe high numbers of longitudinally cut bacterial cells and revealed a new antibacterial mechanism of tetracycline, which is independent from its ability to inhibit the ribosome. The membrane-distorting effect of tetracycline resulted in the complete delocalization of the cell division-regulatory protein couple MinCD, which could explain an earlier observation that certain *B. subtilis* cell division mutants are hypersensitive to tetracycline, a phenomenon that was also independent from the ribosome inhibiting activity of tetracycline (45). Tetracycline also disturbed the localization of the cytoskeletal protein MreB and it is reasonable to assume that more membrane proteins will be affected, which would substantially impact the viability of cells. This additional activity of tetracycline also explains why the ribosomal *tet-4* mutation confers much lower levels of tetracycline resistance (MIC = 16 µg/ml) than the *tetL* resistance cassette that encodes for an efflux pump (MIC = 100 µg/ml). There is still a strong bias against membrane-targeting antibiotics, since they have a reputation to be unspecific and generally toxic. The fact that such an established antibiotic as tetracycline has been ‘secretly’ targeting the bacterial cell membrane for such a long time, underlines that the bacterial cell membrane can be successfully targeted without major side effects on human cells.

Membrane activity has never been shown for tetracycline before, but its analogue anhydrotetracycline has been suspected to target the cell membrane and to cause depolarization. This was based on the fact that anhydrotetracycline causes cell lysis in *E. coli* (46). Our data now show for the first time that anhydrotetracycline does indeed directly affect the cell membrane, however it does not kill by membrane depolarization. Our findings have significant implications for the use of anhydrotetracycline as inducer of gene expression, which is widely advertised to not have antibacterial activity (33). In fact, we have shown that anhydrotetracycline has an even higher antibiotic activity than tetracycline (Supplementary Table 1).

Recently, it was shown that chelocardin, another member of the tetracycline group of antibiotics, inhibits translation but at higher concentrations also causes membrane stress (47). Anhydrotetracycline and chelocardin are often referred to as atypical tetracyclines, which are characterized by being bactericidal. The typical tetracyclines, such as oxytetracycline and tetracycline itself, are bacteriostatic and assumed to only target the ribosome (10). Our study now shows that both groups share membrane distortion as an overarching feature of their antibacterial activity.

How exactly tetracycline affects the cell membrane remains to be investigated. It has been proposed that due to their rather hydrophobic core structure, tetracyclines remain in the cytoplasmic membrane for a relatively long time before they translocate into the cytosol (10, 48). In fact, the clear membrane fluorescence signal observed with both tetracycline and anhydrotetracycline supports this hypothesis. It is reasonable to assume that the same chemical properties that retain these molecules in the membrane also promote bilayer distortion. Tetracycline is a large molecule with a bulky structure, which is likely to disturb the organization of the lipid bilayer. Anhydrotetracycline possesses a methyl group instead of the hydroxyl group, which stimulates interaction with the hydrocarbon core of the lipid bilayer (49). This may explain why anhydrotetracycline has a more severe effect on membrane organization.

From oxytetracycline, which was the first tetracycline to become commercially available in 1950, to doxycycline, which is one of the most commonly prescribed antibiotic drugs today, tetracyclines are widely used in human and veterinary medicine. Despite this heavy use, target-based resistance mutations against tetracycline occur slowly, which has been attributed to the fact that ribosomes are encoded by multiple genes (50). However, resistance against other ribosome inhibitors like streptomycin is frequently observed (51). Therefore, an alternative explanation for the low resistance development against tetracyclines could be that they have a second target, the cell membrane, for which it is generally difficult to obtain suppressor mutations (52). Developing tetracyclines with enhanced membrane effects could be a desirable strategy to combat bacterial infections, since membrane-active bactericidal compounds are often also effective against persister cells, which are an increasing problem in the clinic (53–55). Finally, our results underscore the emerging realization that multi-target antibiotics are most successful in clinical use (50).

## Materials and Methods

### Antibiotics

Gramicidin, vancomycin, ampicillin, nitrofurantoin, tetracycline, anhydrotetracycline, kanamycin, and chloramphenicol were purchased from Sigma-Aldrich in the highest available purity. Daptomycin was purchased from Abcam. MP196 was synthesized by solid-phase synthesis as described previously (56). Gramicidin, nitrofurantoin, anhydrotetracycline, and MP196 were dissolved in sterile DMSO. Vancomycin, ampicillin, kanamycin, and daptomycin were dissolved in sterile water. Tetracycline and chloramphenicol were dissolved in ethanol.

### Strain construction

Strains used in this study are listed in Supplementary Table 2. pAPNC-213-kan-based (57) plasmid pMW4 for integration of *sepF* into the *aprE* locus in the *B. subtilis* genome was constructed by Gibson assembly (58) using the primer pairs MW135 and MW139 for amplifying the pAPNC-213-kan vector backbone and MW140 and MW141 for amplifying the *sepF* gene. pMW4 was transformed into *B. subtilis* 168 using a standard starvation protocol (59). Deletion of the *sepF* gene was accomplished by transforming the resulting *B. subtilis* strain with chromosomal DNA isolated from YK204 (*sepF::spc*) (60). For construction of *B. subtilis* TNVS205 (*aprE::cat-Pspac-mcherry-mreB*) the *mreB* gene was amplified using the primer pair TerS397/TerS400 and the plasmid pAPNC213-cat (61) was amplified with the primer pairs TerS398/TerS337 and TerS338/135. The resulting PCR products were subjected to a three-fragment Gibson assembly reaction resulting in plasmid pTNV86. Transformation of pTNV86 into *B. subtilis* 168 resulted in TNVS205. See Supplementary Table 3 for primer sequences.

### Minimal inhibitory concentration (MIC)

Minimal inhibitory concentrations were determined in a serial dilution assay as described in (62). Briefly, lysogeny broth (LB) was supplemented with different antibiotic concentrations and inoculated with 5×10^5^ CFU/ml of *B. subtilis* 168 (63). Cells were grown at 37 °C under steady agitation for 16 h. The lowest antibiotic concentration inhibiting visible bacterial growth was defined as MIC. The MIC of daptomycin was tested in presence of 1.25 mM CaCl_2_.

### Growth experiments

*B. subtilis* 168 was aerobically grown in LB. Overnight cultures were diluted to an OD_600_ of 0.05 and allowed to grow until an OD_600_ of 0.4 prior to addition of antibiotics at different MIC multiples. Growth was monitored for 8 h using a Biotek Synergy MX plate reader. Concentrations leading to a reduced growth rate without causing massive cell lysis within the first 30 min of antibiotic exposure were chosen for electron microscopy. Daptomycin requires the presence of 1.25 mM CaCl_2_. Addition of CaCl_2_ did not affect growth of *B. subtilis*.

### Growth conditions for TEM experiments

*B. subtilis* 168 and *E. coli* MG1655 were grown in LB. *B. subtilis* MW17 was grown in the presence of 50 µg/ml spectinomycin and 0.5 mM IPTG overnight and diluted 1:100 into antibiotic-free medium containing 0.5 mM IPTG for the embedding experiment. *M. bovis* Bacillus Calmette Guérin (BCG) Tice (64) was grown in 7H9 medium (Difco) supplemented with Middlebrook albumin/dextrose/catalase supplement (BD Biosciences), and 0.05% Tween 80. When *M. bovis* BCG was to be observed without detergent treatment, cultures were washed and resuspended in fresh medium without Tween two days prior to the embedding procedure. *A. laidlawii* PG-8A was grown in modified PPLO medium (1.41% PPLO broth (BD Biosciences), 0.15% TC Yeastolate (Difco), 1.4% glucose, 20% horse serum, 1000 U/ml penicillin G (Sigma-Aldrich)). All cultures were maintained at 37° C under continuous shaking. After reaching mid-logarithmic growth phase, 50 µl of cells were withdrawn, pelleted by centrifugation (16,000x g, 2 min), and resuspended in 5 µl medium. For antibiotic treatment, *B. subtilis* 168 was aerobically grown in LB to an OD_600_ of 0.3 and subsequently treated with either 32 µg/ml valinomycin, 1 µg/ml vancomycin, 1 µg/ml ampicillin, 16 µg/ml MP196, 0.5 µg/ml daptomycin, 32 µg/ml nitrofurantoin, 2 µg/ml tetracycline, 2 µg/ml anhydrotetracycline, or left untreated as control. After 30 min of antibiotic treatment, 50-150 µl of sample were pelleted by centrifugation (16,000x g, 2 min), and resuspended in 5-15 µl fresh LB. Higher cell densities resulted in less effective alignment of cells in the final sections.

### Flat embedding of bacteria on agarose

5-15 µl of the cell suspensions were spotted on 0.25 mm thick 1.5% agarose films (thickness was controlled using Gene Frame AB0576, ThermoFischer Scientific, see Supplementary Movie 1 for preparation). Excess liquid medium was allowed to evaporate under a slight air flow in a clean air bench. A 10 µl drop of sample created an area large enough to produce at least five individual blocks for sectioning. Volumes less than 5 µl still resulted in one or two blocks, making it well possible to work with initial culture volumes of less than 50 µl. For our antibiotic study we chose to spot 10 µl to have more material in case the antibiotic-treated cells would attach less efficiently to the agarose layer. For *M. tuberculosis*, which in general did not attach very well to agarose, we used 15 µl while for the other non-antibiotic treated bacterial samples 5 µl were sufficient. Spotting volumes higher than 15 µl, using higher concentrated samples, or multiple spotting on the same agarose patch did not further increase the number of longitudinally cut cells and in fact compromised their alignment. Agarose patches were transferred to aluminum dishes and kept free-floating (not sticking to the bottom of the dish to allow optimal diffusion of solutions into the agarose film) in the respective solutions with the cell samples facing upwards during all fixation, washing, and dehydration steps. Mounted cells were fixed in 5% glutaraldehyde (Merck) in 0.1 M cacodylate (Sigma-Aldrich) buffer (pH 7.4) for 20 min. Samples were subsequently washed three times with 0.1 M cacodylate, pH 7.4 for 5 min each, followed by incubation in 1% OsO_4_ (EMS) / 1% KRu(III)(CN)_6_ (Sigma Aldrich) for 30 min. Samples were then washed three times with ultrapure water for 5 min each. Dehydration was performed in an incubation series with rising ethanol (Merck) concentrations as follows: 5 min 30% ethanol, 5 min 50% ethanol, 2x 15 min 70% ethanol, 1 h 80% ethanol, 15 min 90% ethanol, 15 min 96% ethanol, 15 min 100% ethanol, 30 min 100% ethanol, water-free. Water-free ethanol was prepared by adding 2 ml acidulated 2,2-dimethoxypropan (two drops of 37% HCl ad 100 ml 2,2-dimethoxypropan (Sigma Aldrich)) to 100 ml ethanol absolute (Merck). Cells were then incubated for 30 min in a 1:1 mixture of EPON and propylene oxide (EMS), followed by 30 min incubation in 2:1 EPON / propylene oxide. All fixation, washing, and dehydration solutions were gently and slowly added starting from the side of the agarose patch in order not to wash off the spotted cells. Agarose patches were transferred to fresh aluminum dishes and covered with fresh EPON. Samples were left at room temperature overnight and then incubated at 65 °C for at least 36 h. EPON was prepared by mixing 48 g glycid ether (Serva) with 32 g dodecenylsuccinic anhydride (Serva) and 20 g methyl nadic anhydride (Serva). Components were mixed for 10 min prior to addition of four times 650 µl benzyldimethylamine (Serva) and mixed for an additional 15 min. EPON aliquots were kept at -20 °C until use. EPON solutions should not be kept for longer than 1 week prior to embedding to avoid infiltration artifacts. For flat embedding, it turned out that EPON prepared with glycid ether is superior to EPON prepared with EMbed (EMS), since the latter results in less flexible EPON discs, which are more difficult to cut when selecting areas of interest for mounting.

### Classical pellet embedding

For pellet embedding, a 50 ml culture was harvested and the resulting pellet was fixed and dehydrated as above, with the only difference that cell pellets were incubated with the different fixation, washing, and dehydration solutions in glass vials under slow agitation. Fixed and dehydrated cell pellets were embedded in EPON using standard conical tip capsules.

### Flat embedding of bacteria on glass slides

Bacteria were grown and concentrated as described above. Concentrated cell suspensions (10 µl) were spotted on glass cover slips that were coated with a thin carbon film as contrast agent to facilitate correct positioning of the EPON block during sectioning. Cells were mounted as described above and subsequently fixed in 5% glutaraldehyde / 0.1 M cacodylate (pH 7.4) for 20 min. Samples were washed three times with 0.1 M cacodylate (pH 7.4) for 5 min each, followed by incubation in 1% OsO_4_ / 1% KRu(III)(CN)_6_ for 30 min. Since in this preparation procedure only the cells themselves and no agarose films need to be dehydrated, shorter dehydration times are possible. After washing the samples three times with ultrapure water (5 min each), dehydration was performed as follows: 5 min 30% ethanol, 5 min 50% ethanol, 5 min 70% ethanol, 30 min 80% ethanol, 15 min 90% ethanol, 15 min 96% ethanol, 5 min 100% ethanol, 15 min 100% ethanol, water-free, 30 min 50% EPON / 50% water-free ethanol. Slides were then transferred to fresh aluminum dishes and further prepared as described above.

Sandwich embedding of *M. bovis* BCG

While *M. bovis* BCG aligned well on agarose, it was prone to subsequently being washed off the surface, which was effectively prevented by enclosing it in an agarose sandwich. To this end, 10 µl of cells were spotted on an agarose patch as described above. After drying, the sample was covered with a second thin layer of 1.5% agarose. Low melting agarose did not give a stable and flat layer, making standard agarose superior for this method. In order not to induce a heat shock, the agarose solution was allowed to cool down to ∼50 °C prior to applying it to the sample. The fresh agarose spot was immediately covered by a glass coverslip to produce a thin and flat surface. Immediately placing a small weight (e.g. a half-full 50 ml falcon tube face down) on top of the glass coverslip resulted in a thinner agarose layer, which greatly facilitates dehydration of the samples. After approximately 1 min the weight was removed and the coverslip was gently slid off the agarose, resulting in a flat and stable agarose sandwich (see Supplementary Movie 2 for preparation). The sandwich samples were further processed like normal agarose-embedded samples as described above.

### Electron microscopy

Regions of interest were selected by observing the EPON-embedded bacterial layer under a light microscope prior to mounting on EPON blocks for thin sectioning. Ultrathin sections (∼80 nm) were cut parallel to the bacterial layer, collected on single-slot, Formvar-coated copper grids, and subsequently counterstained with uranyl acetate (Ultrostain I, Laurylab) and lead citrate (Reynolds) in a Leica EM AC20 ultrastainer. Bacteria were imaged using a JEOL 1010 transmission electron microscope at an electron voltage of 60 kV using a side-mounted CCD camera (Modera, EMSIS).

### Further notes about flat embedding

Although flat embedding is straight forward, does not require any further technology or resources, and can be adapted in any laboratory equipped for TEM, two points have to be taken into consideration when applying this technique. Firstly, embedding on agarose requires careful dehydration, since residual water will lead to infiltration artifacts that either jeopardize ultrathin sectioning or, if sectioning is still possible, appear as strong background in the final sections. Insufficient dehydration might also lead to crooking of the agarose patch after overnight incubation with the EPON resin, defying the purpose of flat embedding. Therefore, dehydration steps should not be shortened and the agarose layer must be thin, especially for the sandwich approach. In this respect, flat embedding on carbon-coated glass surfaces is clearly at advantage, since in this case only the cells need to be dehydrated. Secondly, since all cells are aligned in one plane in both agarose and glass methods, ultrathin sectioning requires an experienced person in order to hit the resin block at perfectly perpendicularly angle to the cells. While finding the right plane is easier in glass-embedded samples due to the carbon film, it also requires higher precision and care, since the almost perfect alignment of cells limits the tolerance for failed sectioning attempts.

### Fluorescence light microscopy

All strains were aerobically grown in LB until an OD_600_ of 0.4 prior to antibiotic treatment. For Nile red staining *B. subtilis* 168 was treated with 2 µg/ml tetracycline, anhydrotetracycline, 15 µg/ml chloramphenicol, or 3 µg/ml kanamycin for 30 min followed by membrane staining with 0.5 µg/ml Nile red for 1 min. For DAPI staining *B. subtilis* 168 was treated with 32 µg/ml nitrofurantoin for 5, 15, 30, or 60 min, respectively, followed by staining of the chromosome with 1 µg/ml DAPI for 1 min. *B. subtilis* LB318 (168 *amyE::spc mgfp-minD aprE::cat mcherry-minC*) (37) was grown in the presence of 0.1% xylose to induce expression of *mgfp-minD* and 0.1 mM IPTG to induce expression of *mcherry-minC*. TNVS205 (168 *aprE::cat mcherry-mreB*) was grown in the presence of 0.3 mM IPTG to induce expression of *mcherry-mreB*. *B. subtilis* LB318 and TNVS205 were treated with 2 µg/ml tetracycline, 2 µg/ml anhydrotetracycline, or 1 µg/ml gramicidin, respectively. Note that LB318 carries both a chloramphenicol and a kanamycin resistance cassette and TNVS205 carries only chloramphenicol resistance. Concentrations of chloramphenicol and kanamycin were 15 and 3 µg/ml, respectively, for non-resistant strains, and 20 and 10 µg/ml for strains carrying the respective resistance marker(s), which corresponds to double the selection concentration. Samples were observed under the microscope after 5 and 30 min of antibiotic treatment. Staining with DiSC(3)5 was carried out as described by te Winkel *et al*. (11) followed by treatment with 2 µg/ml tetracycline, 2 µg/ml anhydrotetracycline, or 1 µg/ml gramicidin, respectively. Samples were examined after 5 and 30 min of antibiotic staining. DiIC12 staining was carried out as described in Müller *et al*. (40). All microscopy samples were spotted on a thin film of 1.2% agarose (11) and examined with a Nikon Eclipse Ti equipped with a CFI Plan Apochromat DM 100x oil objective, an Intensilight HG 130 W lamp, a C11440-22CU Hamamatsu ORCA camera, and NIS elements software. Images were analyzed using ImageJ (National Institutes of Health).

### Structured Illumination Microscopy (SIM)

Samples were prepared as for fluorescence light microscopy. Cover slips were coated with poly-dopamine to reduce background fluorescence by preventing binding of the membrane dye to the glass surface (11). Cells were imaged with a Nikon Eclipse Ti N-SIM E microscope setup equipped with a CFI SR Apochromat TIRF 100x oil objective (NA1.49), a LU-N3-SIM laser unit, an Orca-Flash 4.0 sCMOS camera (Hamamatsu Photonics K.K.), and NIS elements Ar software. Images were analyzed using ImageJ (National Institutes of Health).

### Spectroscopic membrane potential measurements

Cells were cultured as for microscopy experiments and transferred to a pre-warmed 96-well plate after reaching an OD_600_ of 0.4. DiSC(3)5 measurements were carried out as described by te Winkel *et al.* (11). Cells were treated with 2 µg/ml tetracycline, 2 µg/ml anhydrotetracycline, and 1 µg/ml gramicidin and measurements were taken every 30 sec over a total of 30 min. Kanamycin (3 µg/ml) and chloramphenicol (15 µg/ml) were also tested but had no effect on the membrane potential (data not shown). All antibiotics were tested for an effect on DiSC(3)5 fluorescence in solution to control for interference with the dye but no change in DiSC(3)5 fluorescence was observed (data not shown).

## Acknowledgments

We would like to thank Bruce Koppen for assistance with MIC and growth experiments, Tjalling Siersma for assistance with DiSC(3)5 measurements, Laura Bohorquez for constructing LB318, and Terrens Saaki for constructing TNVS205. MP196 was kindly supplied by Nils Metzler-Nolte, Ruhr University Bochum. This work was financially supported by the Netherlands Organization for Scientific Research (NWO, http://nwo.nl/en, STW-Vici 12128 to LWH). MW was supported by a postdoc stipend from the Amsterdam Infection and Immunity Institute. BW was supported by a PhD fellowship of the China Scholarship Council. Electron microscopy was performed at the VU/VUMC EM facility, supported by the Netherlands Organization for Scientific Research (NWO, middelgroot 91111009).

## Supplementary Information

**Supplementary Table 1:** Minimal inhibitory concentrations of antimicrobial compounds against *B. subtilis* 168.

| compound | MIC (µg/ml) |
| --- | --- |
| valinomycin | 16 |
| vancomycin | 0.5 |
| ampicillin | 0.5 |
| daptomycin | 1 |
| MP196 | 32 |
| nitrofurantoin | 8 |
| tetracycline | 8 |
| anhydrotetracycline | 4 |

**Supplementary Table 2:** Bacterial strains used in this study.

| strain name | relevant genotype | induction | reference |
| --- | --- | --- | --- |
| *B. subtilis* 168 | - |  | (63) |
| *B. subtilis* MW17 | *sepF::spc aprE::kan Pspac-sepF* | 1 mM IPTG | this study |
| *B. subtilis* LB318 | *amyE::spc mgfp-minD aprE::cat*<br>*mcherry-minC* | 0.1 mM IPTG,<br>0.1% xylose | (37) |
| *B. subtilis* TNVS205 | *aprE::cat-Pspac-mcherry-mreB* | 0.3 mM IPTG | this study |
| *B. subtilis* PG112 | *tet-4* | - | (45) |
| *B. subtilis* SG82 | *lacA::tet* | - | (45) |
| *E. coli* MG1655 | - | - | (65) |
| *M. bovis* BCG Tice | - | - | (64) |
| *A. laidlawii* PG-8A | - | - | (66) |

**Supplementary Table 3:**
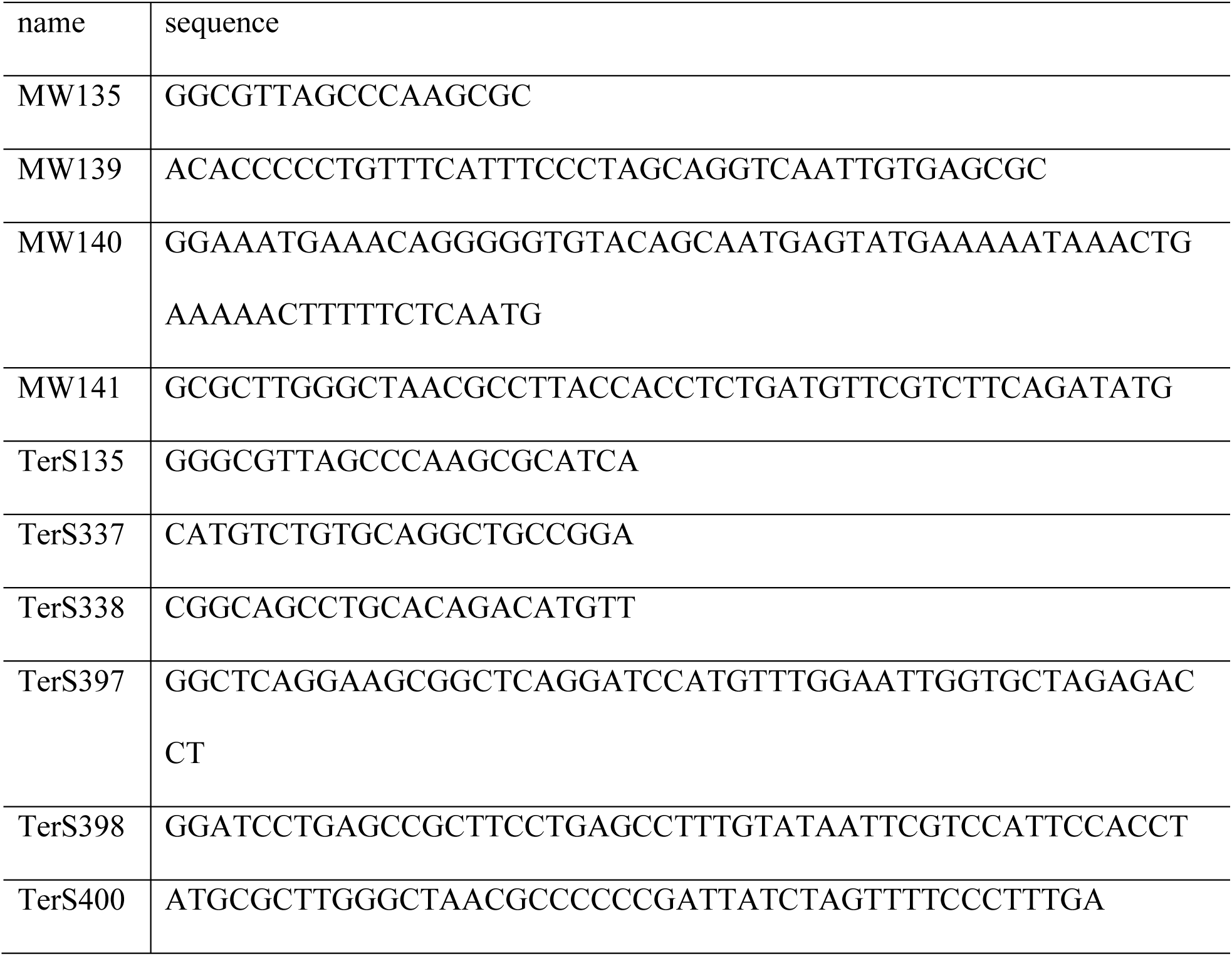
Primer sequences.

**Supplementary Figure 1:**
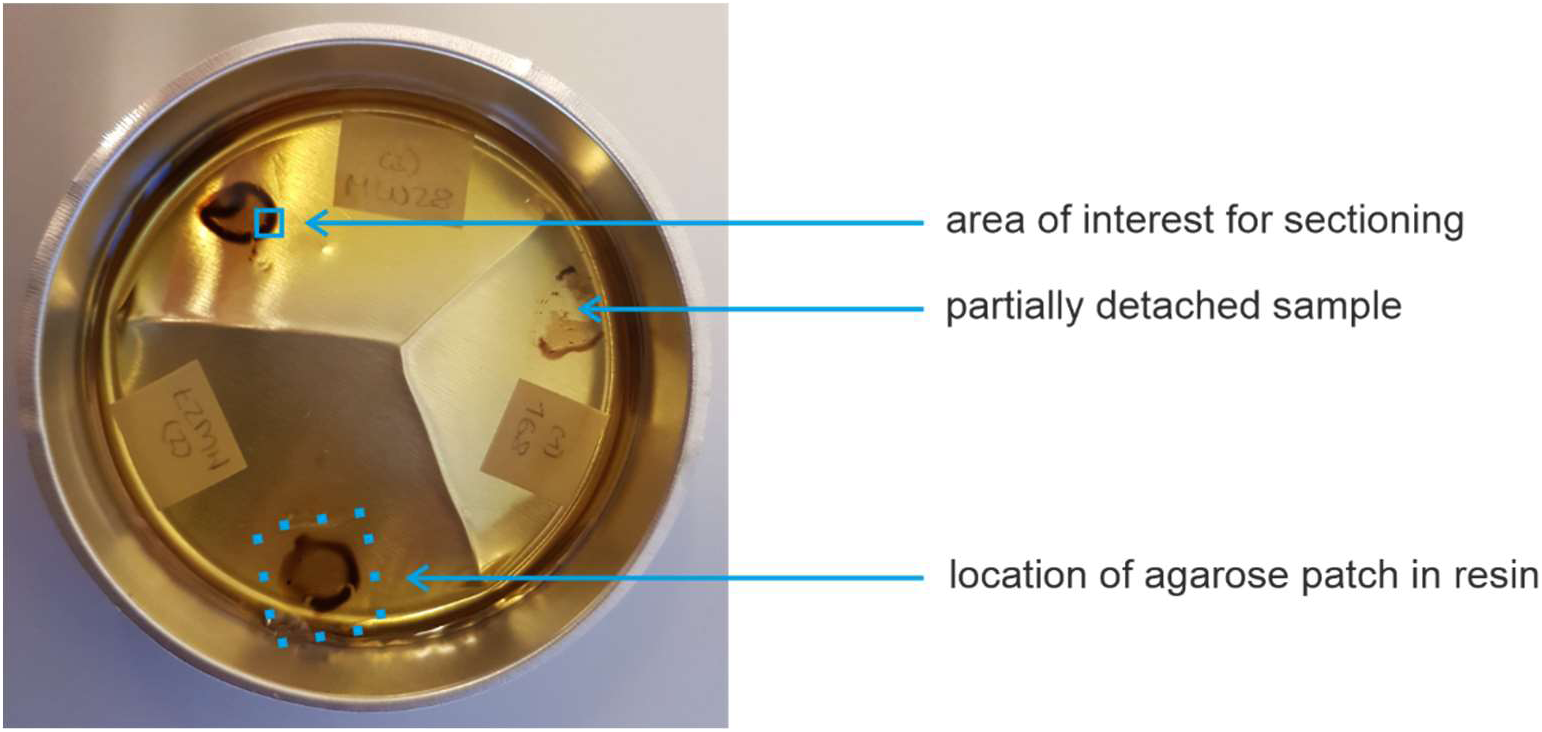
Example of a polymerized EPON disc after flat embedding. The samples shown are from 150 µl logarithmically growing *B. subtilis* cultures that were pelleted and resuspended in 15 µl LB. The whole 15 µl cell suspension was spread on an agarose patch and embedded according to the single agarose layer protocol. The aluminium dish can be removed from the EPON disc and an area of interest can be cut out with a hot scalpel and mounted on a conventional EPON block for ultrathin sectioning. From these sample volumes, a minimum of 5 sectioning blocks can be prepared. Nicely aligned cells can typically be found in the middle of the sample or close to the dark halo. Within the halo itself, cells were more prone to overlap with each other, resulting in less complete longitudinally cut cells in the final sections. However, for low concentrated samples, certain mutants, and partially lysing cultures we made the experience that the dark halo gives better sections than the center of the spot. Therefore, we typically select an area of interest that contains both areas. Disc diameter 7 cm.

**Supplementary Figure 2:**
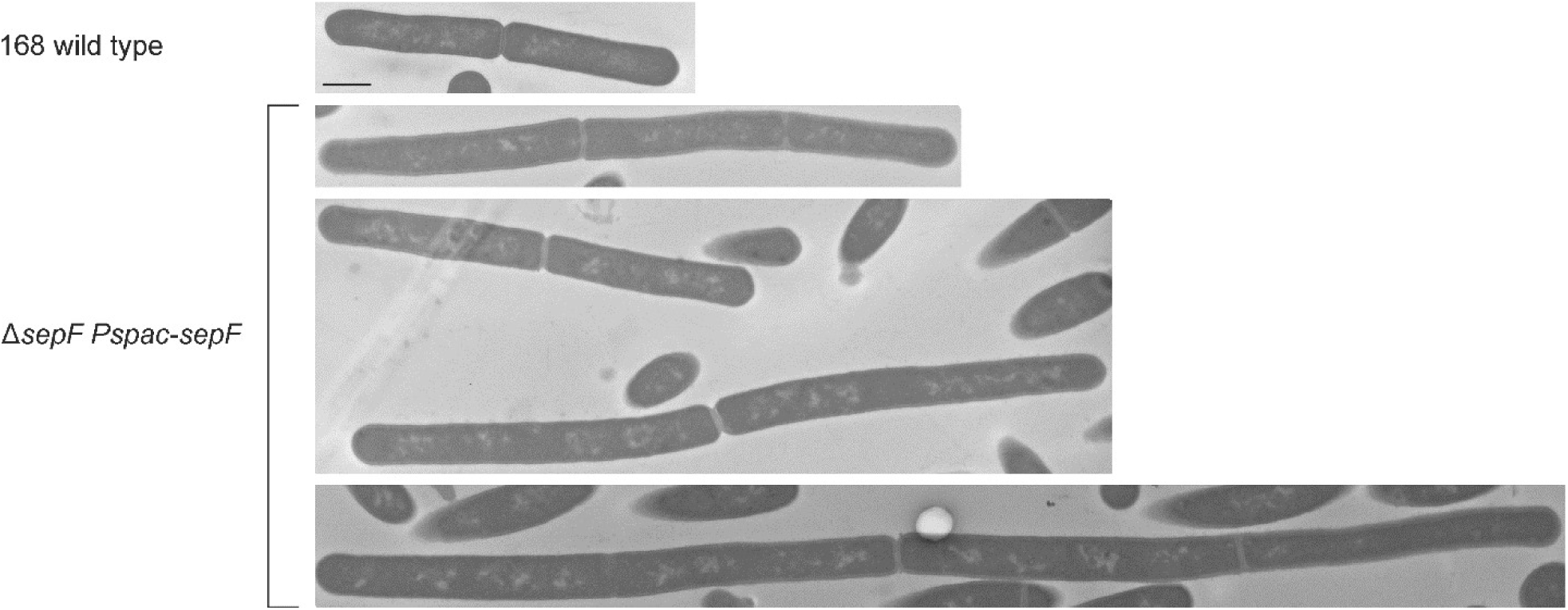
Flat embedding of filamentous cells. Overexpression of the cell division protein SepF inhibits cell division by preventing septum formation (67). Strain MW17 (*B. subtilis* 168 *sepF::spc aprE::kan Pspac-sepF*) carries an IPTG-inducible copy of the *sepF* gene in the ectopic *aprE* locus. Induction with 0.5 mM IPTG results in elongated cells (67). Scale bar 1 µm.

**Supplementary Figure 3:**
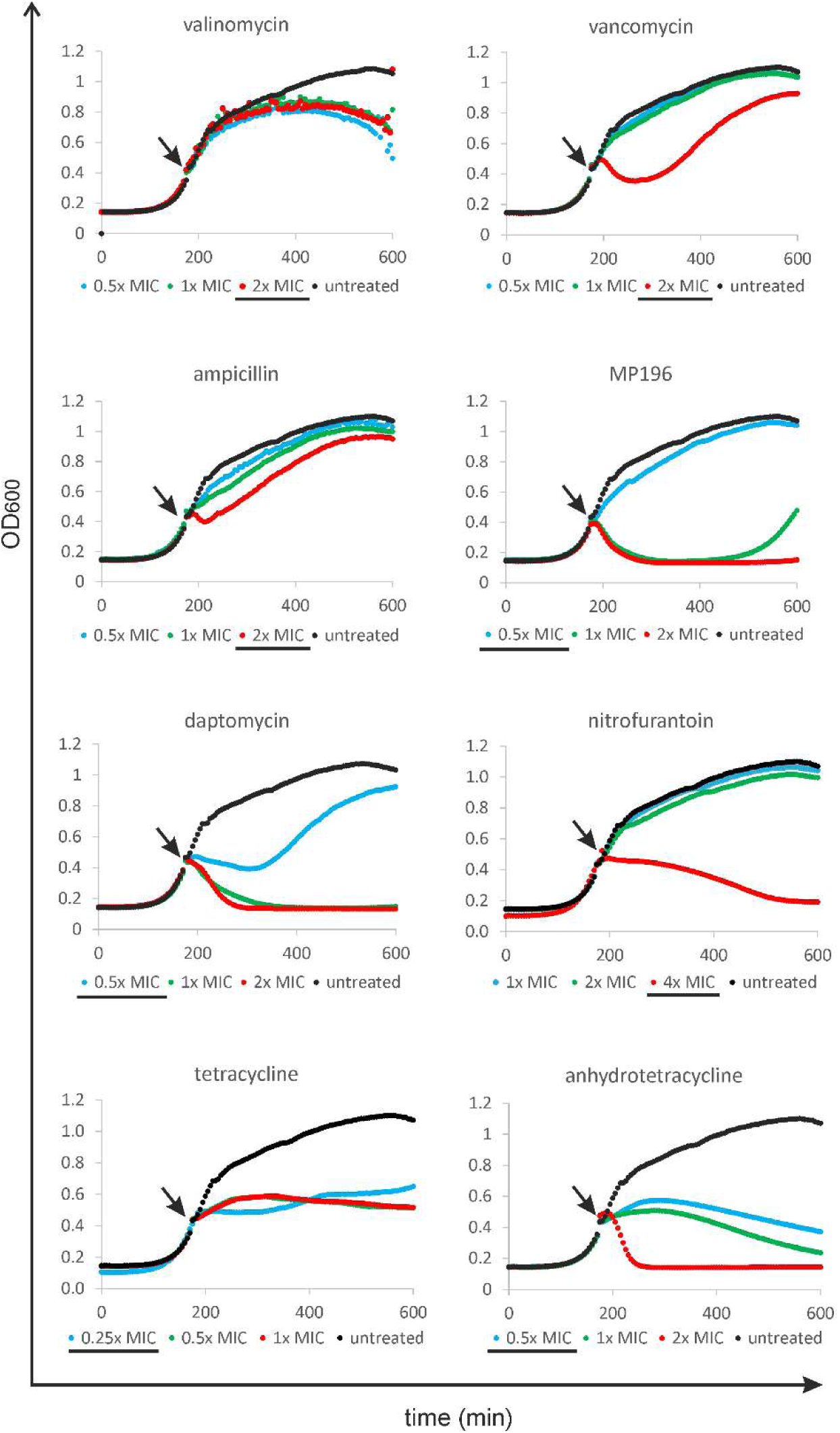
Growth of *B. subtilis* 168 after treatment with different antibiotic concentrations. Arrows indicate time points of antibiotic addition. Concentrations used for further experiments are underlined.

**Supplementary Figure 4:**
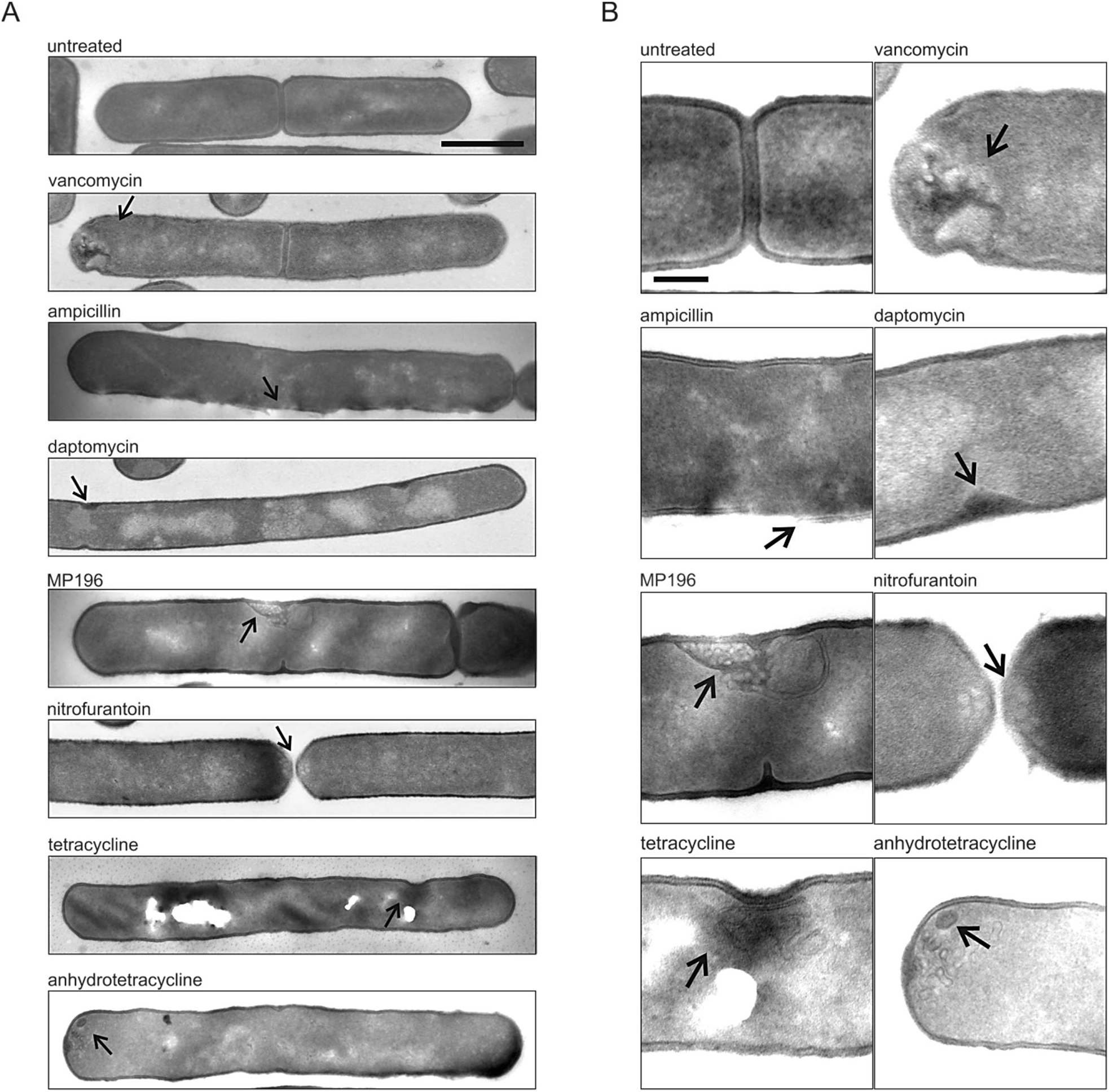
Electron micrographs **(A)** and detail images **(B)** of *B. subtilis* cells treated with different antibiotics for 30 min. Antibiotic-induced lesions are indicated by arrows. We mainly chose antibiotics that target the cell wall and should cause clearly visible cell wall defects. Vancomycin binds to the cell wall precursor molecule lipid II and thus inhibits cell wall synthesis. Cells treated with this antibiotic show clear cell wall lesions. Ampicillin inhibits transpeptidation of peptidoglycan polymers (21), causing cell wall thinning and ultimately cell lysis (68). Accordingly, ampicillin-treated cells displayed partly disintegrated cell walls. Daptomycin was recently shown to hamper cell wall synthesis by targeting membrane microdomains that harbor the cell wall synthetic machinery, causing them to accumulate into lipid II-enriched foci (40, 69). In line, daptomycin-treated cells showed aberrant local cell wall thickening. The antimicrobial peptide MP196 caused intracellular cell wall structures and membrane vesicles, reflecting its dual mechanism of targeting membrane function and cell wall synthesis (6). Nitrofurantoin is thought to kill cells by an unspecific mechanism involving oxidative damage (23). Cells treated with this antibiotic lacked a nucleoid and showed membrane aberrations, which is consistent with oxidative damage to these cellular structures. Tetracycline inhibits the bacterial ribosome (10). Surprisingly, we consistently observed membrane lesions in tetracycline-treated cells. Anhydrotetracycline, an analogue of tetracycline, which is thought to rather target the cell membrane than the ribosome (32), caused similar lesions. Scale bars 1 µm (A) and 250 µm (B).

**Supplementary Figure 5:**
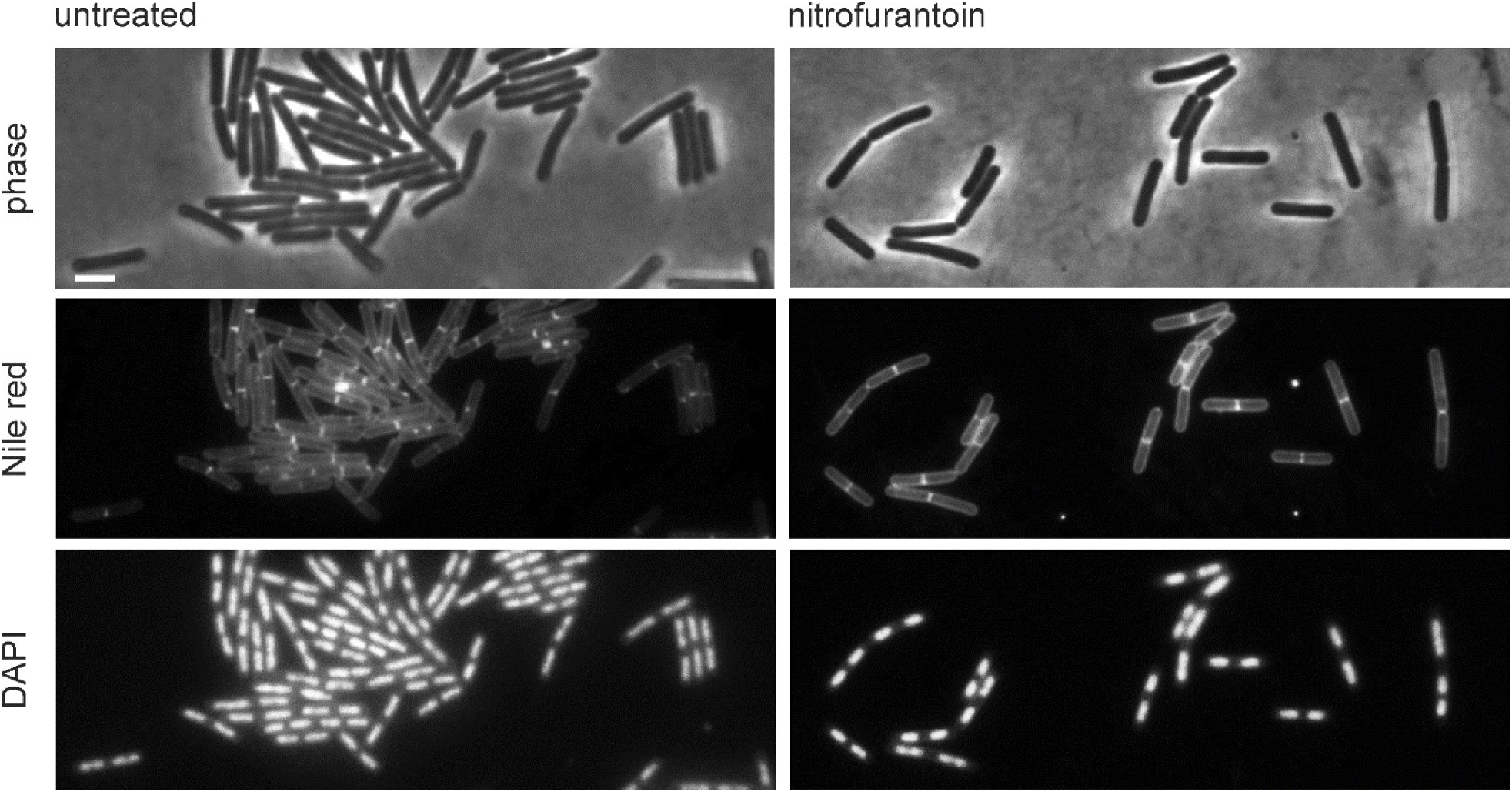
Effects of 5 min treatment with nitrofurantoin on the nucleoid. Scale bar 2 µm.

**Supplementary Figure 6:**
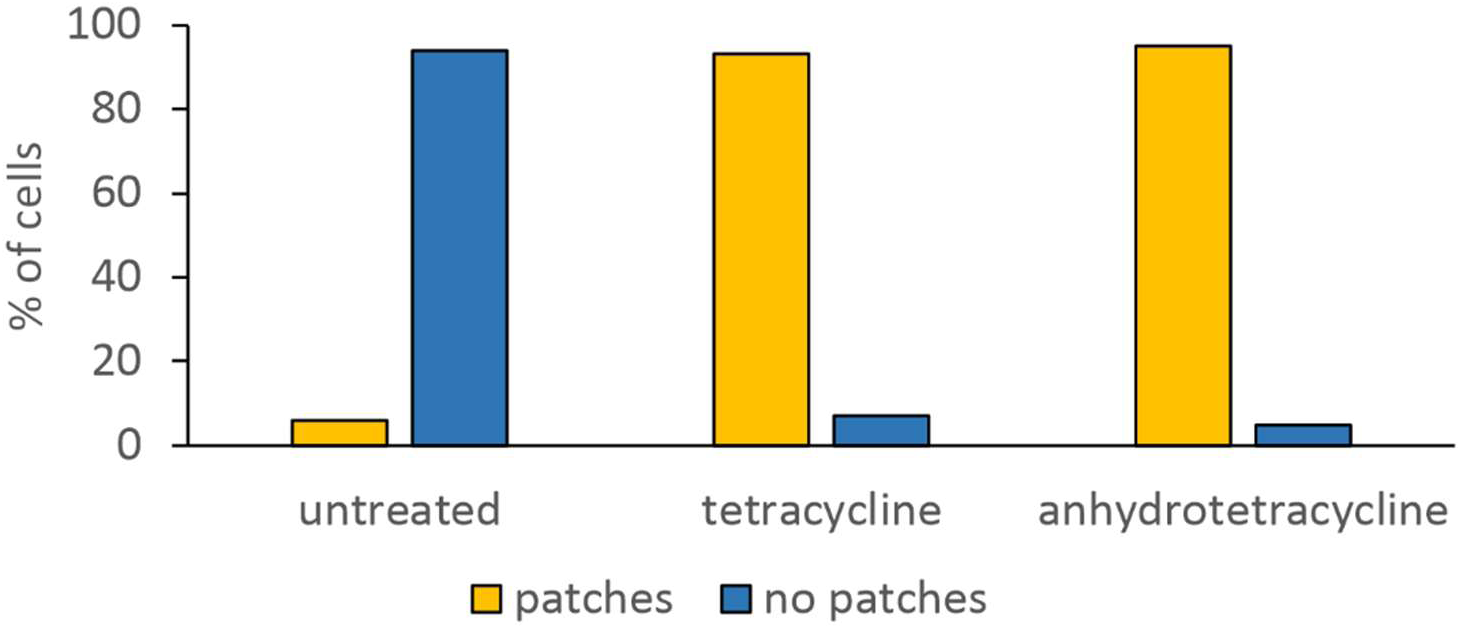
Quantification of fluorescence microscopy images. Cells were inspected for the presence of Nile red-stained fluorescent membrane patches. A minimum of 100 cells were evaluated per condition.

**Supplementary Figure 7:**
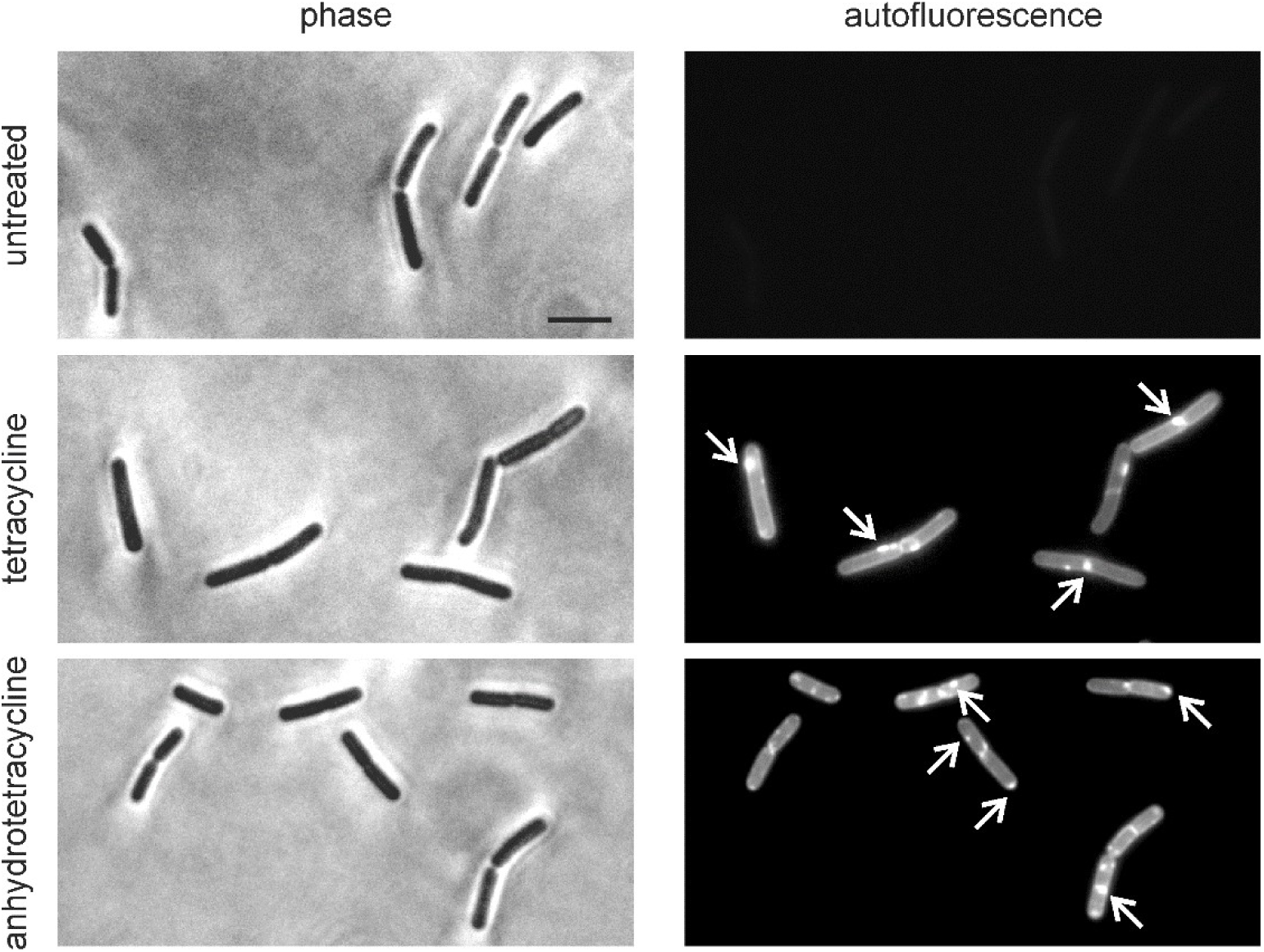
Localization of tetracycline and anhydrotetracycline in *B. subtilis* 168. Green autofluorescence of the tetracyclines allows label-free localization of these antibiotics in living cells. Arrows indicate some sites of compound accumulations in the cell membrane. Scale bar 2 µm.

**Supplementary Figure 8:**
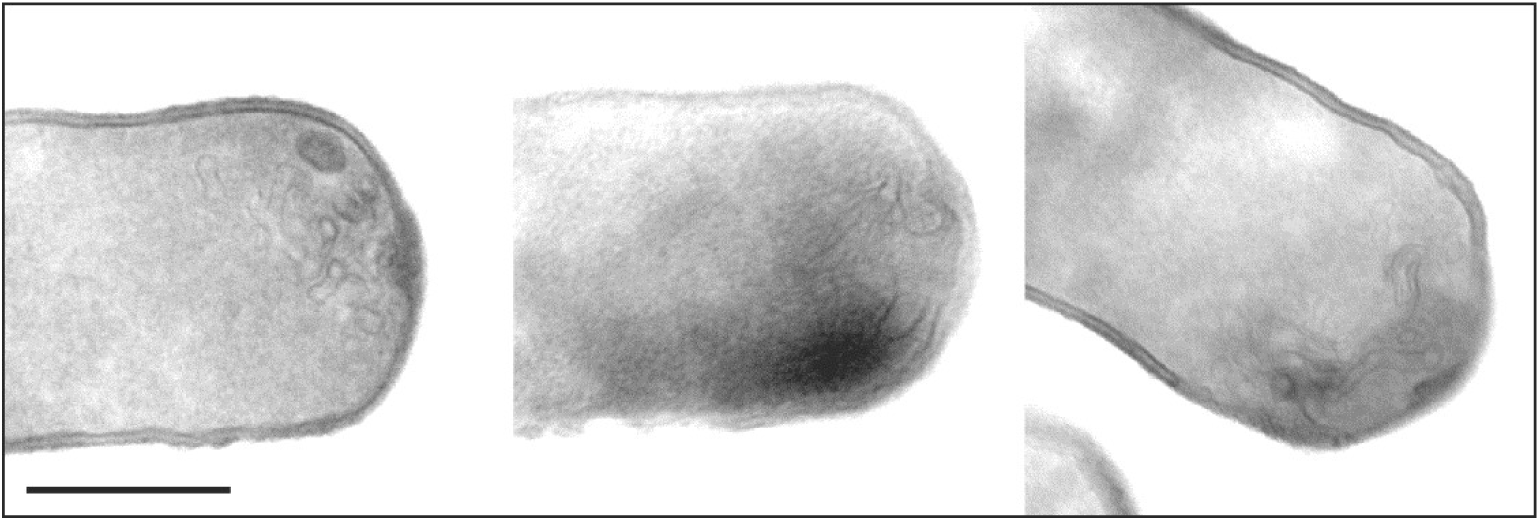
Exemplary lesions caused by anhydrotetracycline. Scale bar 500 nm.

**Supplementary Figure 9:**
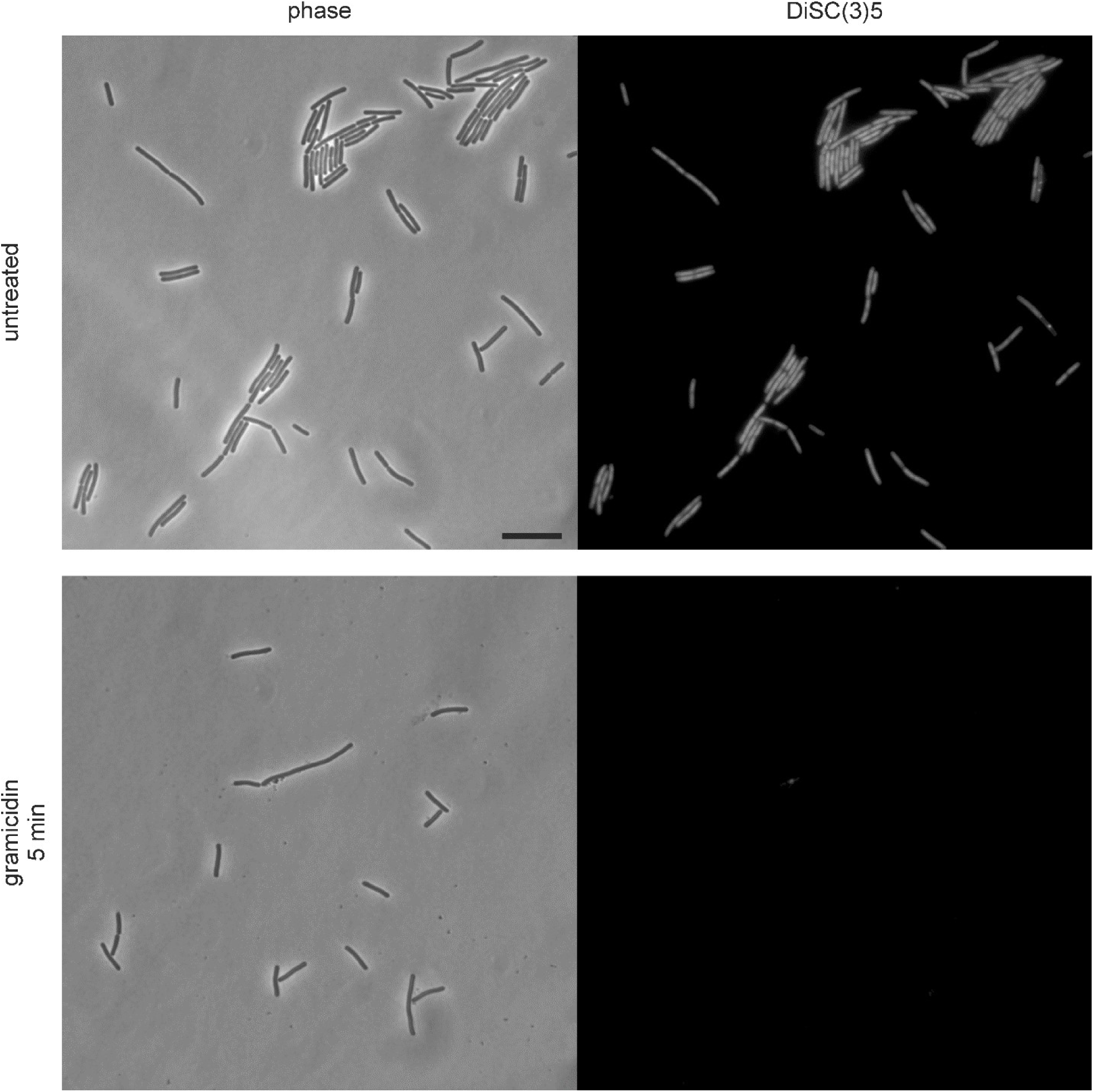
DiSC(3)5 staining of untreated cells (negative control) and cells treated with gramicidin (1 µg/ml, positive control). A fluorescence signal indicates the presence of a membrane potential (negative control: untreated cells). Depolarization leads to release of the dye from the cells and a diminished fluorescence signal in the cells (positive control: gramicidin). All fluorescence pictures in Supplementary Figure 9–11 have been recorded with the same exposure time and were adjusted with the same brightness and contrast settings. Scale bar 10 µm.

**Supplementary Figure 10:**
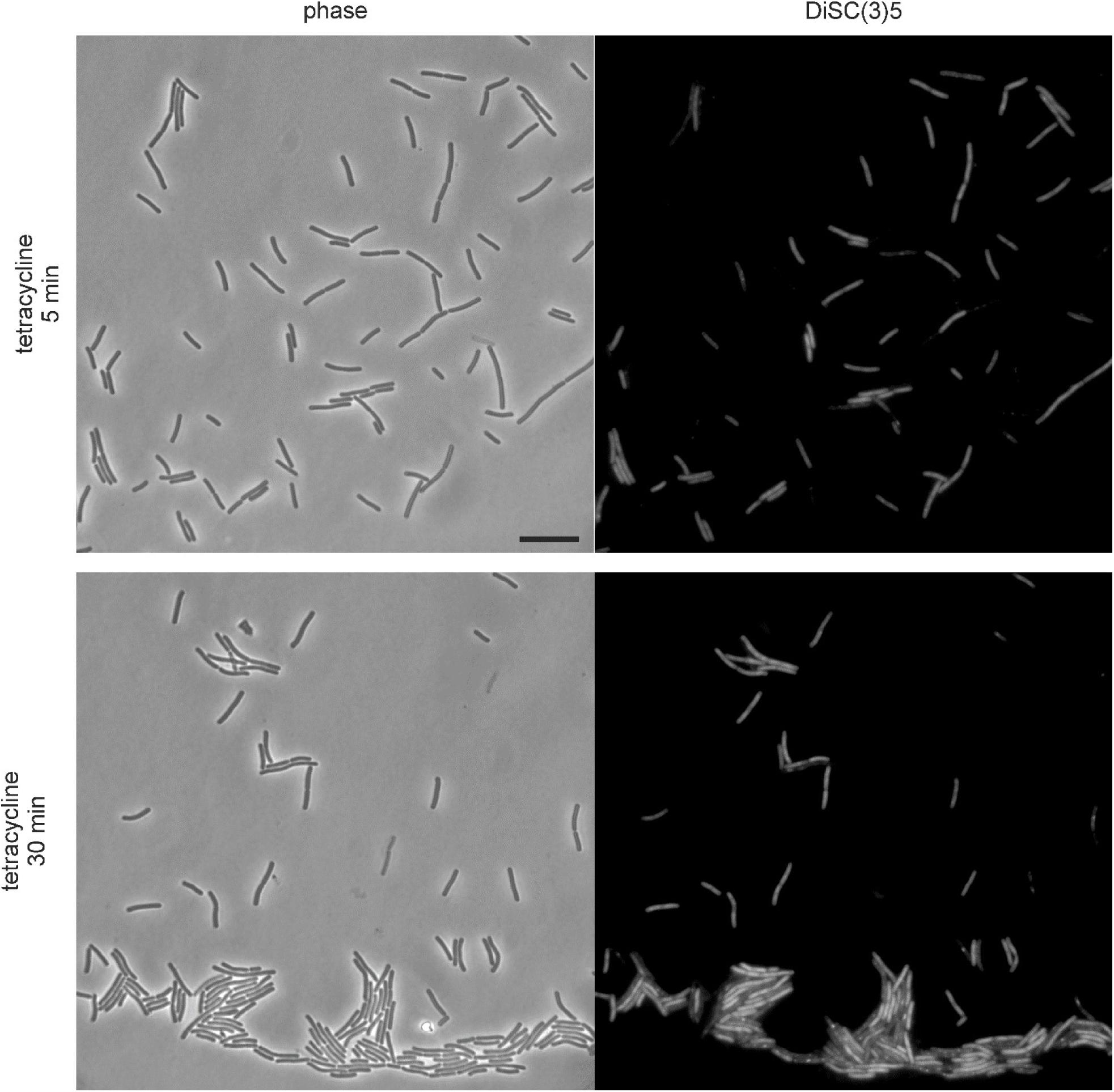
DiSC(3)5 staining of cells treated with tetracycline (2 µg/ml). Note the heterogeneity in the DiSC(3)5 staining. A fluorescence signal indicates the presence of a membrane potential. Depolarization leads to release of the dye from the cells and a diminished fluorescence signal in the cells. All fluorescence pictures in Supplementary Figure 9–11 have been recorded with the same exposure time and were adjusted with the same brightness and contrast settings. Scale bar 10 µm.

**Supplementary Figure 11:**
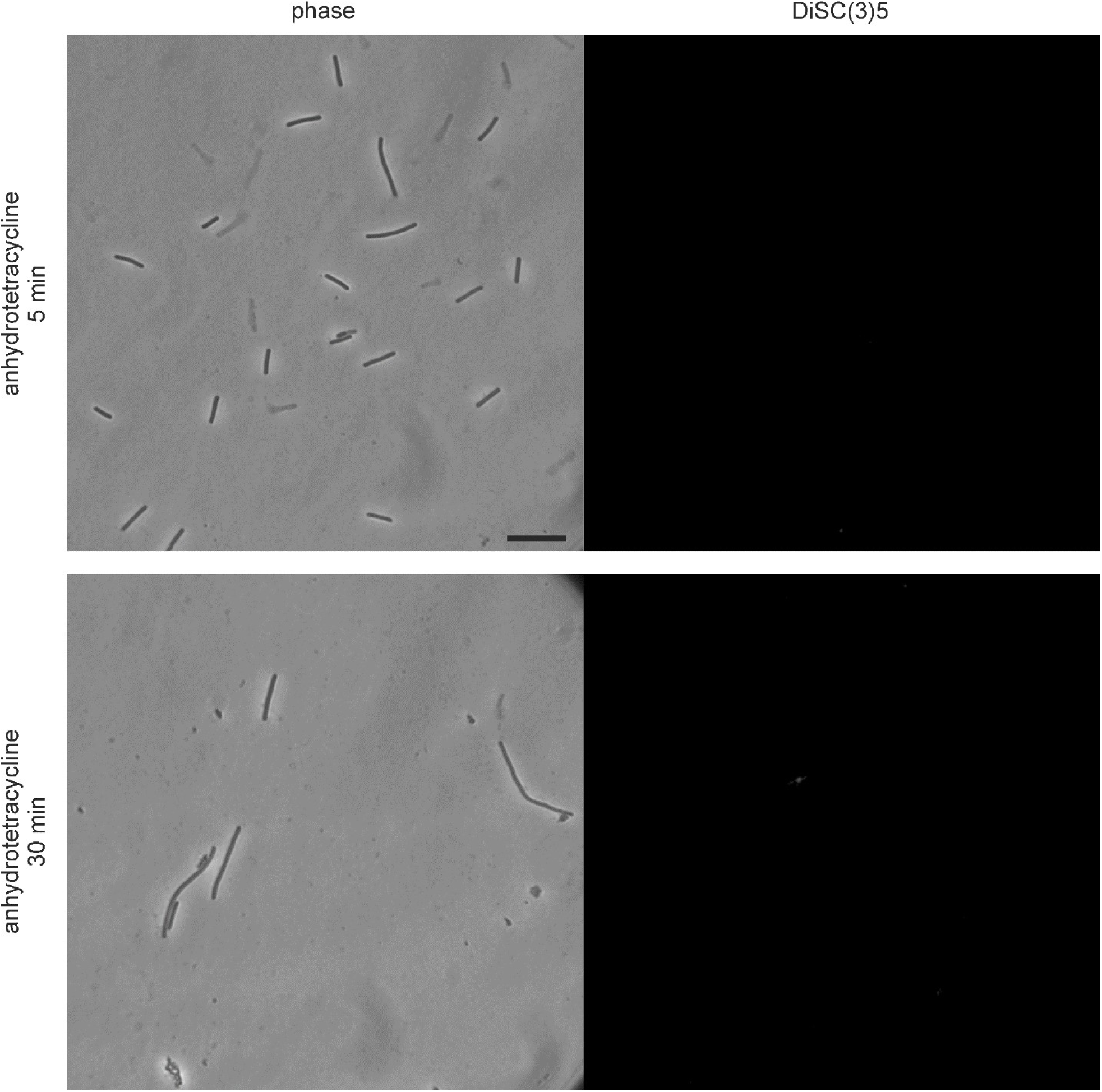
DiSC(3)5 staining of cells treated with anhydrotetracycline (2 µg/ml). A fluorescence signal indicates the presence of a membrane potential. Depolarization leads to release of the dye from the cells and a diminished fluorescence signal in the cells. All fluorescence pictures in Supplementary Figure 9–11 have been recorded with the same exposure time and were adjusted with the same brightness and contrast settings. Scale bar 10 µm.

**Supplementary Figure 12:**
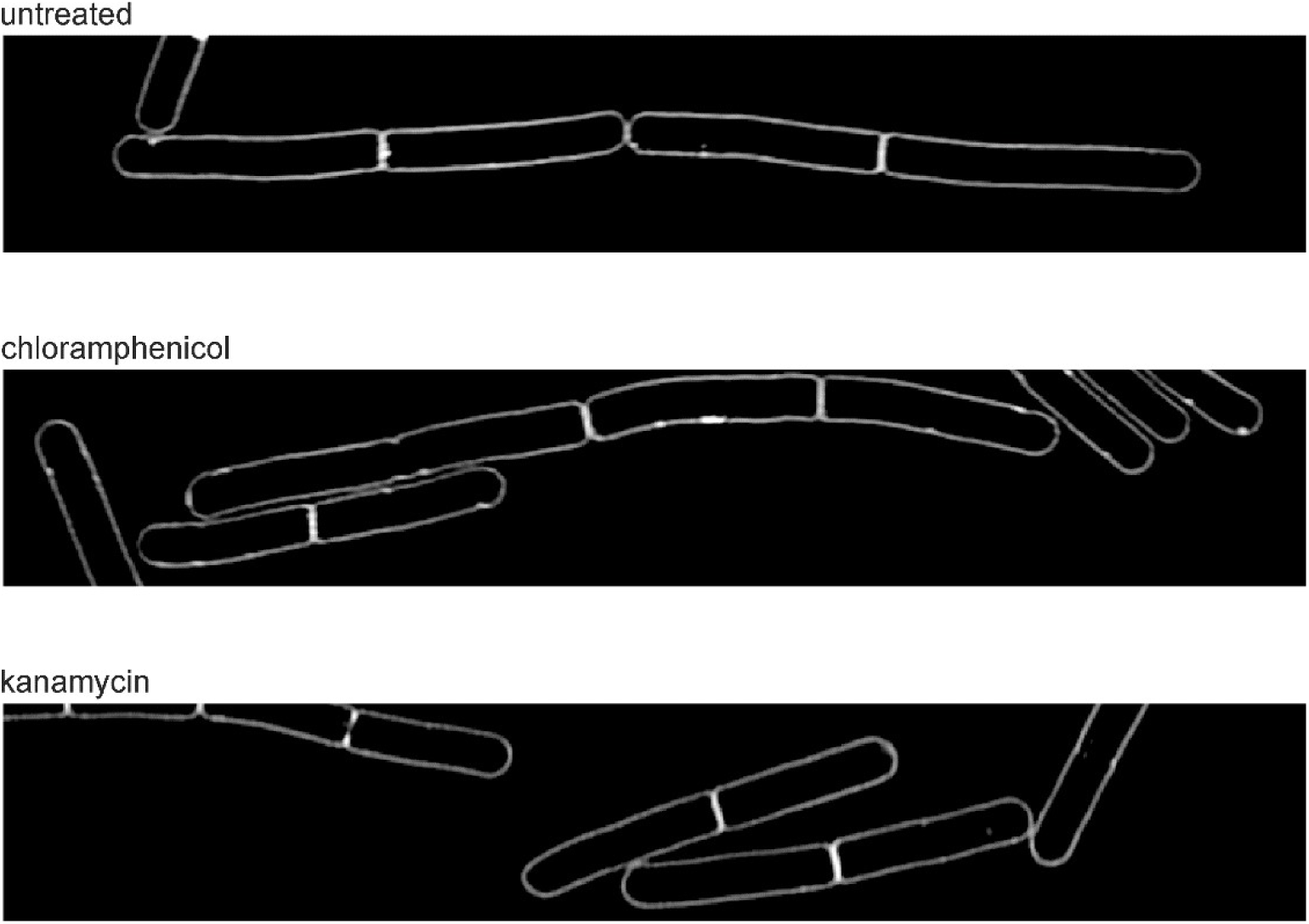
Inhibition of translation does not cause membrane aberrations. Logarithmically growing *B. subtilis* 168 cultures were treated with 15 µg/ml chloramphenicol or 3 µg/ml kanamycin for 20 min, stained with Nile red, and examined by SIM microscopy.

**Supplementary Figure 13:**
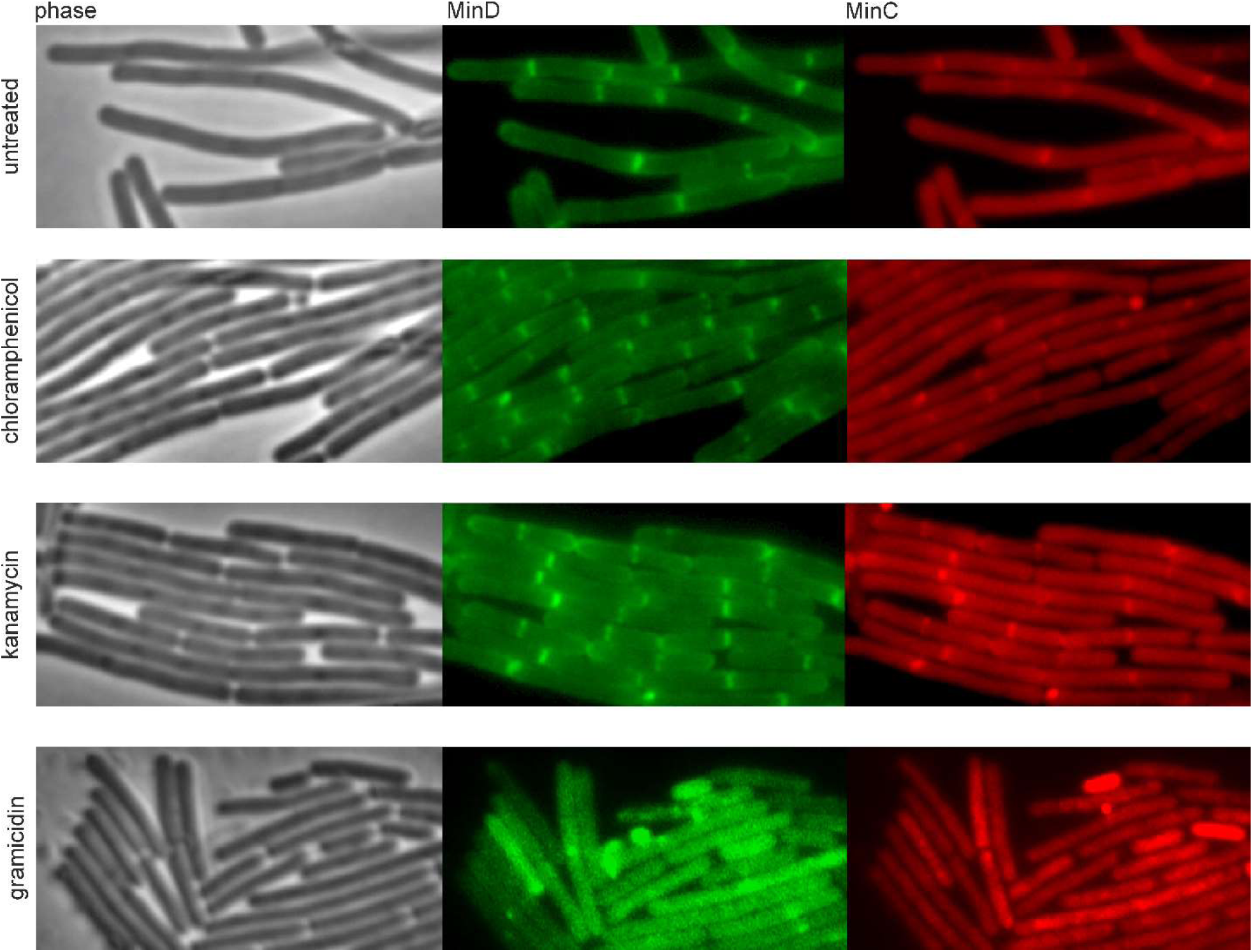
Inhibition of translation does not cause delocalization of the membrane potential-dependent membrane proteins MinD and MinC. *B. subtilis* LB318, expressing GFP-MinD and mCherry-MinC, was treated with 20 µg/ml chloramphenicol, 10 µg/ml kanamycin, or 1 µg/ml gramicidin for 20 min prior to microscopy. Note that LB318 carries both a chloramphenicol and kanamycin resistance cassette. Therefore, twice the concentrations used for antibiotic selection was chosen for microscopy.

**Supplementary Figure 14:**
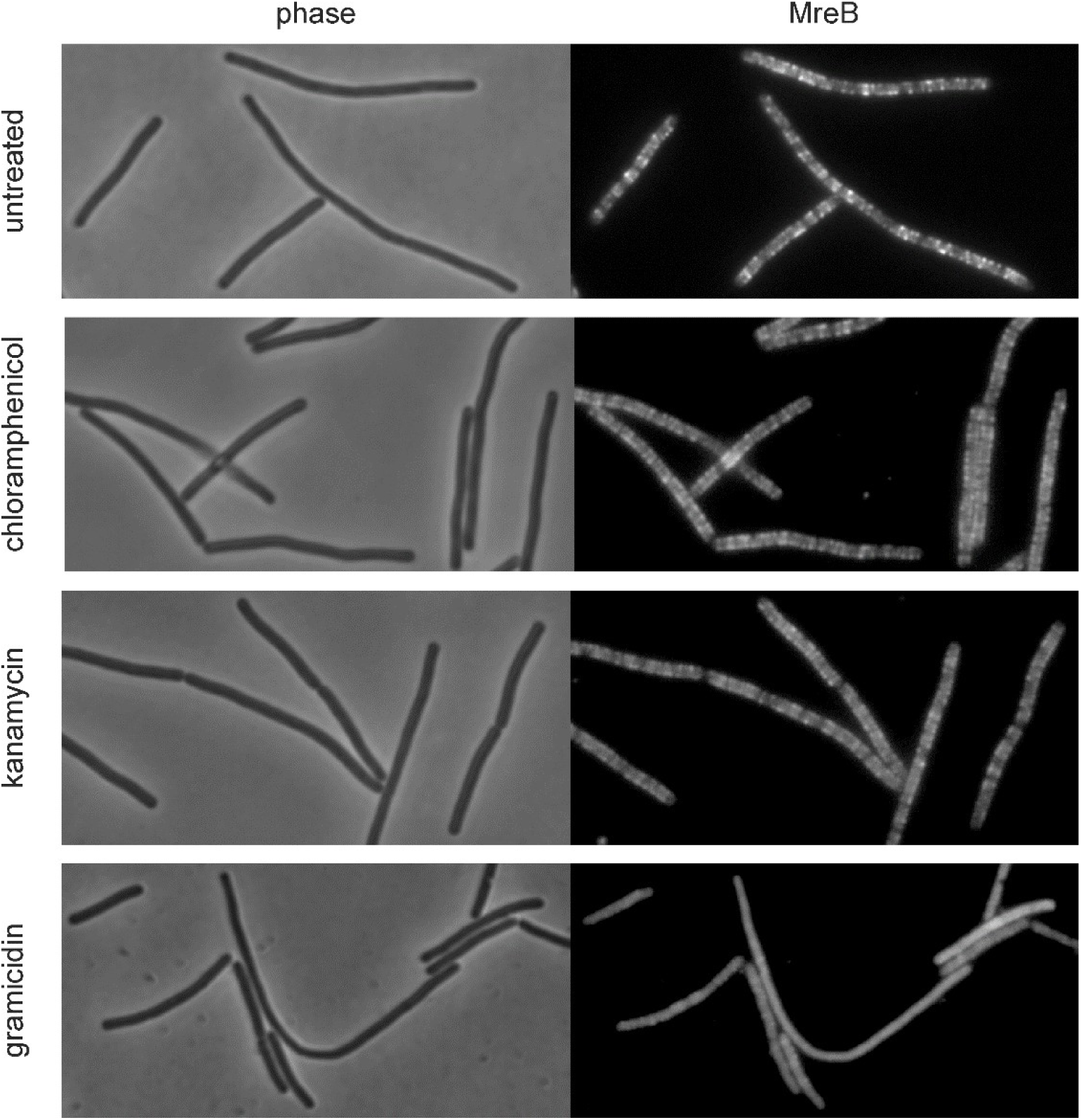
Inhibition of translation does not cause delocalization of MreB. *B. subtilis* TNVS205, expressing mCherry-MreB, was treated with 20 µg/ml chloramphenicol, 3 µg/ml kanamycin, or 1 µg/ml gramicidin for 20 min prior to microscopy. Note that TNVS205 carries a chloramphenicol resistance cassette. Therefore, twice the concentration used for antibiotic selection was chosen for microscopy.

**Supplementary Figure 15:**
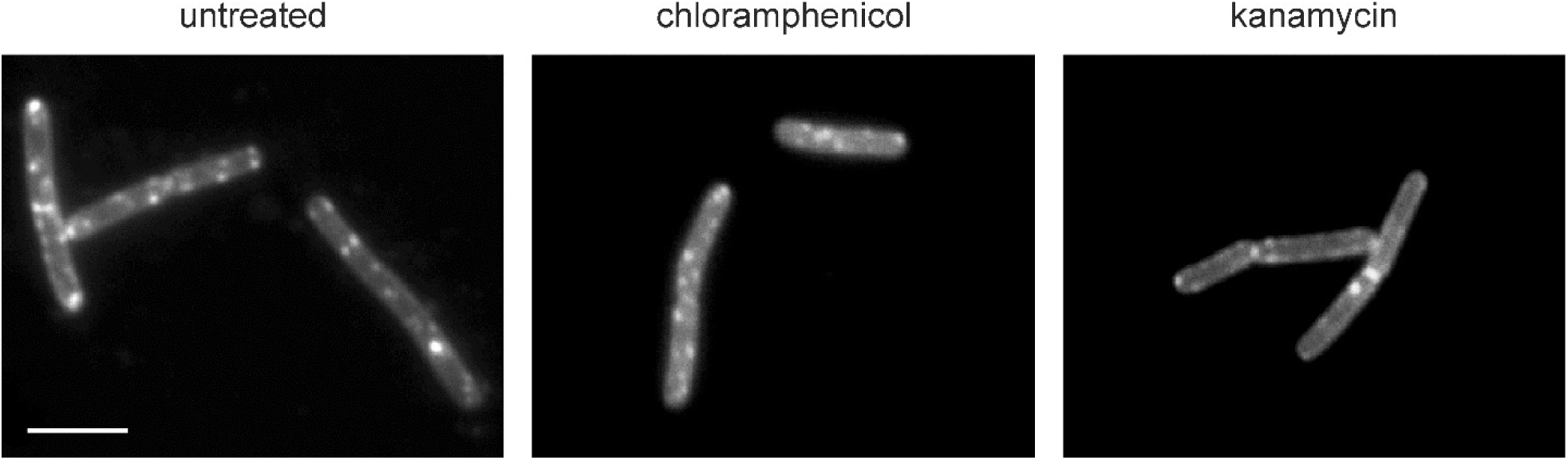
Inhibition of translation does not diminish fluid membrane domains. *B. subtilis* 168 was treated with 15 µg/ml chloramphenicol or 3 µg/ml kanamycin 30 min prior to microscopy. Small effects are expected since RIFs depend on the growth phase (29) and a reduced growth rate caused by antibiotic treatment is likely to have secondary effects on RIFs. In line, RIFs were less clear after 30 min treatment with chloramphenicol and kanamycin compared to the untreated control. However, clustering or diminishing of RIFs was not observed. Scale bar 2 µm.

**Supplementary Figure 16:**
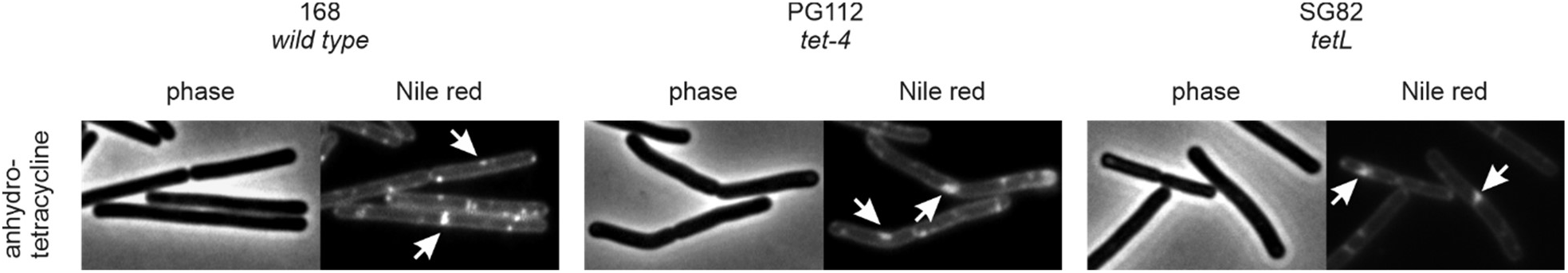
Effect of anhydrotetracycline on tetracycline-resistant strains. Anhydrotetracycline is insensitive to both *tet-4* and *tetL* resistance mechanisms (MIC 1 µg/ml for both PG112 and SG82).

